# TREX2 component PCID2 scaffolds alternative SAC3-based subcomplexes with distinct RNA processing and export function

**DOI:** 10.64898/2026.04.13.716336

**Authors:** Vasilisa Aksenova, Elizabeth Giordano, Caroline Esnault-Petrov, Alexei Arnaoutov, Mary Dasso

## Abstract

The TREX2 complex bridges transcription and RNA export. Its subunits show differences in expression, localization, and dynamics, suggesting distinct cellular roles. To understand the roles of individual TREX2 components, we characterized their interactomes. We identified novel, evolutionarily conserved SAC3(PCI-fold)-based subcomplexes with its PCID2 subunit. PCID2 acts as a scaffold for mutually exclusive yet structurally related subcomplexes with GANP, LENG8, and SAC3D1. These subcomplexes have alternative localization at the nuclear envelope, nuclear speckles and cytosol. LENG8 localizes in nuclear speckles and interacts extensively with the mRNA processing factors. LENG8 depletion alters mRNA processing and polyadenylation site usage. LENG8 thus acts upstream of the canonical TREX2 complex, in which PCID2 cooperates with GANP in mRNA export. Together, our findings reveal that TREX2 is not a uniform complex but a modular system, in which TREX2 subunits can assemble into functionally distinct subcomplexes through interacting partners that define their specificity and alternative functions.

**HIGHLIGHTS:**

- TREX2 subunits play distinct roles in RNA retention and exhibit subunit-specific preference for interactions with different protein partners
- PCID2 forms mutually exclusive subcomplexes with GANP, LENG8, and SAC3D1
- GANP, LENG8, and SAC3D1 alter PCID2 intracellular localization
- LENG8 is a nuclear speckle protein that modulates alternative mRNA processing

## INTRODUCTION

Isolation of RNA transcription within the nucleus, away from cytoplasmic translation, allows eukaryotic RNAs to undergo extensive modifications before becoming functional and being exported to the cytoplasm. For mRNAs, maturation typically involves 5’ capping, co-transcriptional or post-transcriptional splicing, and 3’ end formation, followed by the formation of export-competent ribonucleotide-protein (RNP) particles (1). This complex set of events requires numerous protein complexes that may reside at distinct sites within the nucleus, and mRNAs are subject to strict quality control prior to export. Among the key protein complexes involved in mRNA maturation and export, the TRanscription and EXport 2 (TREX2) complex plays a central role in bridging the transcription, mRNA processing and export machinery (2). The TREX2 complex is well conserved between budding yeast and mammals. Mammalian TREX2 components include GANP (S. cerevisiae homologue: Sac3), PCID2 (Thp1), two copies of ENY2 (Sus1), DSS1 (Sem1), and either CETN2 or CETN3 (Cdc31) proteins (3).

The TREX2 complex and its subunits have been well characterized structurally (3). In particular, GANP is the human orthologue of yeast Sac3, sharing a conserved domain called the SAC3 homology domain as well as a CID domain (Cdc31 interaction domain) that are found across eukaryotic species (Figure 1a). Human GANP also possesses an FG-rich NUP homology domain, and a MCM3AP domain (MCM3-acetylating protein) that are not well conserved in fungi (4). GANP binds to the other TREX2 subunits and serves as a scaffold for complex assembly. The GANP protein shuttles between the nucleoplasm and nuclear envelope, where it is essential for export of mRNA transcripts from the nucleus (3–6). GANP binds to the nuclear pore complex through association to the basket nucleoporin TPR (5) and recruits other TREX2 complex subunits during the process of mRNA export (3, 5). PCID2 is a homolog of the yeast Thp1 protein (7), shuttling between the nucleus and the cytosol (8). In yeast, Thp1 and Sac3 bind each other through their PCI domains, creating a critical surface for binding nucleic acids (9, 10). Similarly, the human Thp1 homolog PCID2 uses its PCI domain to interact with GANP. The yeast Sac3-Thp1 and human GANP-PCID2 complexes are structurally similar to each other (10–14), and are also structural similar to the fungal yCsn12-Thp3 complex (15). Yeast Thp3 has a notable role in regulating transcription and mRNA splicing (15). PCID2 and GANP further interact with DDX39B (also known as UAP56), a multifunctional regulator of nuclear mRNP maturation and export (11, 12, 14).

**Figure 1.**
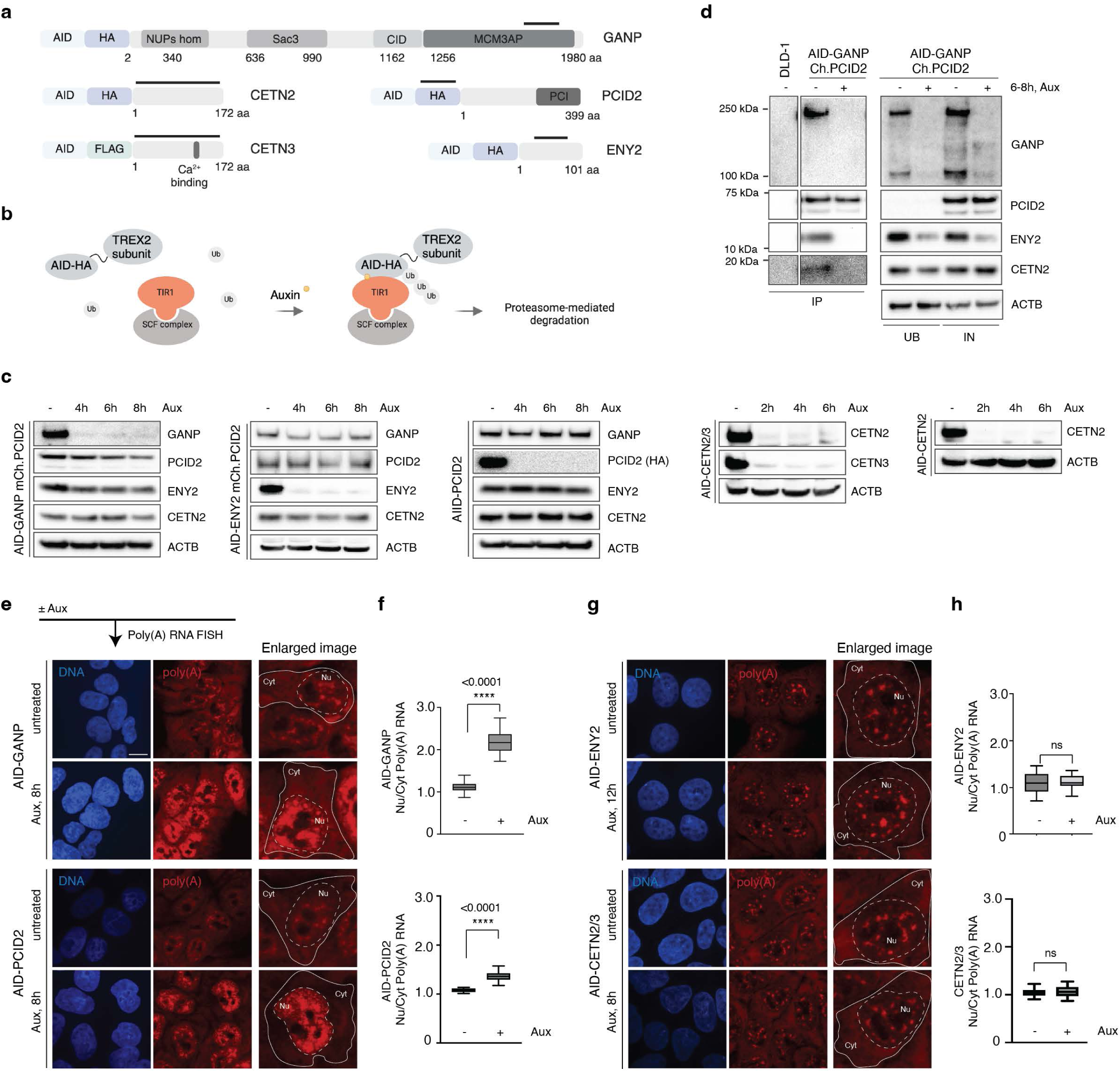
TREX2 subunits differentially contribute to global RNA retention. **a**, A schematic of CRISPR/Cas9-based tagging of TREX2 subunits genes with AID degron and HA or FLAG tag. Each AID cell line is biallelically tagged. Bold lines highlight immunogenic regions within TREX2 subunits or tag used for antibodies production and detection on Western blotting. **b**, A model of auxin-mediated rapid degradation of TREX2 subunits. **c**, Degradation dynamics of GANP, ENY2, PCID2, CETN2-CETN3, and CETN2, and subunits stability after 2-8 h of auxin treatment. **d**, Analysis of TREX2 complex assembly through immunoprecipitation of mCherry-PCID2 from untreated and GANP-depleted cells. AID – Auxin Inducible Degron, HA - human influenza hemagglutinin. **e**, **g**, GANP and PCID2 (e) but not ENY2 or CETNs (g) required for efficient export of poly(A) RNA. **f**, **h**, Quantification of Nu/Cyt Poly(A) RNA distribution.

While TREX2 complex subunits can be biochemically purified together and show similar phenotypes upon siRNA-mediated depletion (4), their functional behavior suggests that they do not operate in perfect tandem. For example, the expression of different subunits varies across tissues (Supplementary Fig.1), their residence times at nuclear pores vary (6) and distinct genetic disorders are linked to their genes (7, 16–21). We thus wished to not only determine how each of these subunits functions in mRNA processing and export, but also whether any of them act outside of the context of the TREX2 complex. To address this question, we developed biochemical methods for the analysis of proteins associated to each of the TREX2 subunits.

Here we describe the identification of interacting partners of TREX2 subunits that can determine their specificity and alternative functions, with a focus on subcomplexes formed by PCID2. Our studies revealed novel and evolutionarily conserved subcomplexes of PCID2 with LENG8 and SAC3D1, two proteins that have SAC3 (PCI-fold) domains similar to GANP. PCID2 binds to all three proteins through their SAC3-homology domains, forming mutually exclusive yet structurally related subcomplexes. Further investigation suggests that each of these PCID2-containing complexes has a pattern of localization and function that are distinct from the others. For LENG8 in particular, our findings indicate that it localizes at nuclear speckles, sites of concentration for proteins involved in mRNA splicing. LENG8 depletion alters patterns of mRNA splice site selection coupled with a shift in alternative polyadenylation site usage. Notably, these findings not only demonstrate that the LENG8 controls important aspects of alternative mRNA processing but also indicate more broadly that distinct interacting partners of TREX2 subunits can diversify the functions of the TREX2 complex for roles outside of mRNA export.

## RESULTS

### TREX2 subunits differentially contribute to RNA retention and transcriptome profiles

We wished to determine the phenotypes associated with loss of individual TREX2 subunits. To do this, we created cell lines wherein each subunit protein was tagged with an Auxin-Induced Degron (AID) tag and either a HA or FLAG epitope, except for DSS1, for which tagging was unsuccessful (Figures 1a and S2, Table S1). Using an auxin-mediated system (Figure 1b), each TREX2 subunit could be depleted within 2-8 h after auxin addition (Figure 1c). After depletion of any individual TREX2 component, we analyzed the stability of non-targeted subunits. Consistent with earlier reports, we found that loss of either GANP or ENY2 impacted each other’s stability (6), whereas the loss of PCID2 did not impact the stability of the other subunits (Figure 1c). As anticipated, we found co-precipitation of all subunits, reflecting their association within the TREX2 complex. Loss of the scaffold subunit GANP resulted in the disassembly of the complex, disrupting interactions between PCID2, ENY2, and CETN2 (Figure 1d).

This system allowed us to track mRNA export and transcriptomic outcomes for each subunit independently. We performed fluorescent in situ hybridization (FISH) against poly(A) RNA to assess bulk RNA export from the nucleus to the cytoplasm. While loss of GANP or PCID2 led to measurable retention of poly(A) mRNA inside the nucleus (Figure 1e-f), depletion of ENY2 or of both CETN2 and CETN3 did not (Figure 1g-h). These results show a strong requirement for GANP and PCID2 in bulk poly(A) RNA trafficking that is not shared with all subunits, arguing that the absence of GANP, PCID2, ENY2 or CETNs does not produce equivalent phenotypes.

We used RNA-seq analysis to further assess how TREX2 subunits shape the cellular transcriptome. We observed that loss of GANP, PCID2, ENY2 and CETNs led to distinct profiles, with significant changes in the abundance of 2480, 1341, 913 and 3 RNAs (differential expression (DE) ≥ 30%, adj. p value <0.05, DESeq2 normalized counts >1), respectively, further indicating that TREX2 subunits are functionally distinct (Figure 2a, b). The transcriptomic changes associated with GANP and PCID2 loss were most closely aligned to each other and to the profile obtained after the loss of the basket nucleoporin TPR (Figure 2a, (5)). By contrast, the loss of ENY2 resulted in a distinct transcriptomic profile that differed substantially from changes after the loss of GANP and PCID2 (Figure 2a, Table S2A), while loss of CETNs caused minimal transcriptomic changes. More transcripts were downregulated than upregulated following the loss of TREX2 subunits (Figure 2c-d). The transcripts downregulated upon GANP and PCID2 depletion tended to be encoded by genes with fewer introns than their upregulated counterparts, resulting in shorter total exon lengths and higher GC content. This trend was reversed in ENY2-dependent transcripts, where downregulated genes exhibited a higher number of introns, had longer exons and lower GC content (Figure 2e). Gene Ontology (GO) analysis revealed that most of the transcripts differentially and specifically impacted upon GANP, PCID2, and ENY2 loss are involved in regulating metabolic processes, including nucleic acid biosynthesis, transcription regulation by polymerase II, RNA metabolism, and RNA processing (Figure 2f-g), which aligns with the expected roles of the TREX2 subunits in bridging transcription and RNA export (3).

**Figure 2.**
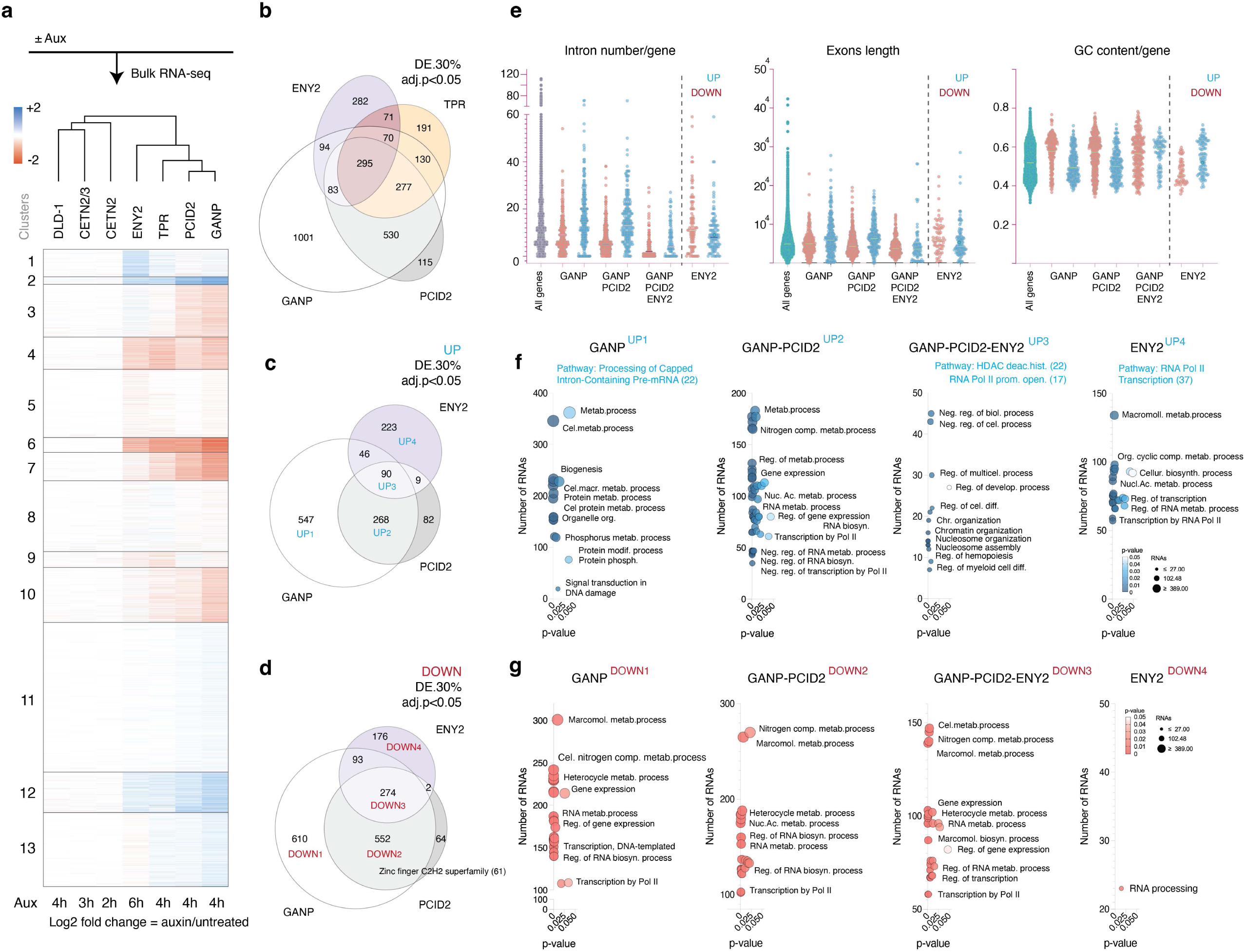
TREX2 subunits do not share similar transcriptomic responses upon their loss. **a,** A heatmap of unsupervised k-means clustering of differentially expressed genes 2-6 h after auxin treatment of cells expressing corresponding AID-tagged CETN2/3, CETN2, ENY2, TPR, PCID2, GANP and auxin-treated DLD-1 cells. **b**, A Venn diagram representing the number of RNAs that showed significant change (both up- and down-regulation) upon ENY2, TPR, GANP, or PCID2 loss. DE log2FC>30%, adj. p-value < 0.05, DESeq2 normalized counts > 1. **c, d,** A Venn diagram showing overlap of upregulated and downregulated transcripts upon GANP, PCID2, and ENY2 loss. **e-g,** Properties (e) and GO terms (f-g) of up-(blue) and down-(red) regulated RNAs affected by GANP, GANP or PCID2, GANP or PCID2 or ENY2, or ENY2 loss. UP1-4 and DOWN1-4 groups on f-g and c-g correspond to each other. Three independent biological replicates were used to perform RNA-Seq of indicated cell lines. DE – Differentially Expressed, UP – upregulated, DOWN – downregulated.

Collectively, both poly(A) RNA retention and RNA-seq analysis demonstrated a closer alignment of PCID2 and GANP, as compared to the impact of ENY2 or CETNs depletion. These findings argue that while the TREX2 subunits work together, they do not function in perfect tandem and potentially have individual roles.

### Identification of binding partners for TREX2 subunits

Given the distinct phenotypes after depletion of individual TREX2 subunits, we examined their association to other proteins outside the TREX2 complex. For this analysis, we created cell lines with mNeonGreen (NG)-tagged GANP, PCID2 and ENY2, as well as cells in which CETN2 and CETN3 were tagged with HA and FLAG epitopes, respectively (Figures 3a and S2). In all cases, the tags were inserted biallelically at the endogenous gene loci. We visualized NG-tagged GANP, PCID2 and ENY2 localizations using live imaging (Figure 3b). As expected, NG-tagged GANP was localized at the nuclear rim, while PCID2 and ENY2 were found at the nuclear rim, in the nucleoplasm, and in intranuclear clusters (6). Immunostaining of HA-CETN2 and FLAG-CETN3 cell lines showed that CETNs were localized within the nucleus or centrosomes (Figure 3b), a pattern that did not reproduce nuclear rim localization observed in other reports (6).

**Figure 3.**
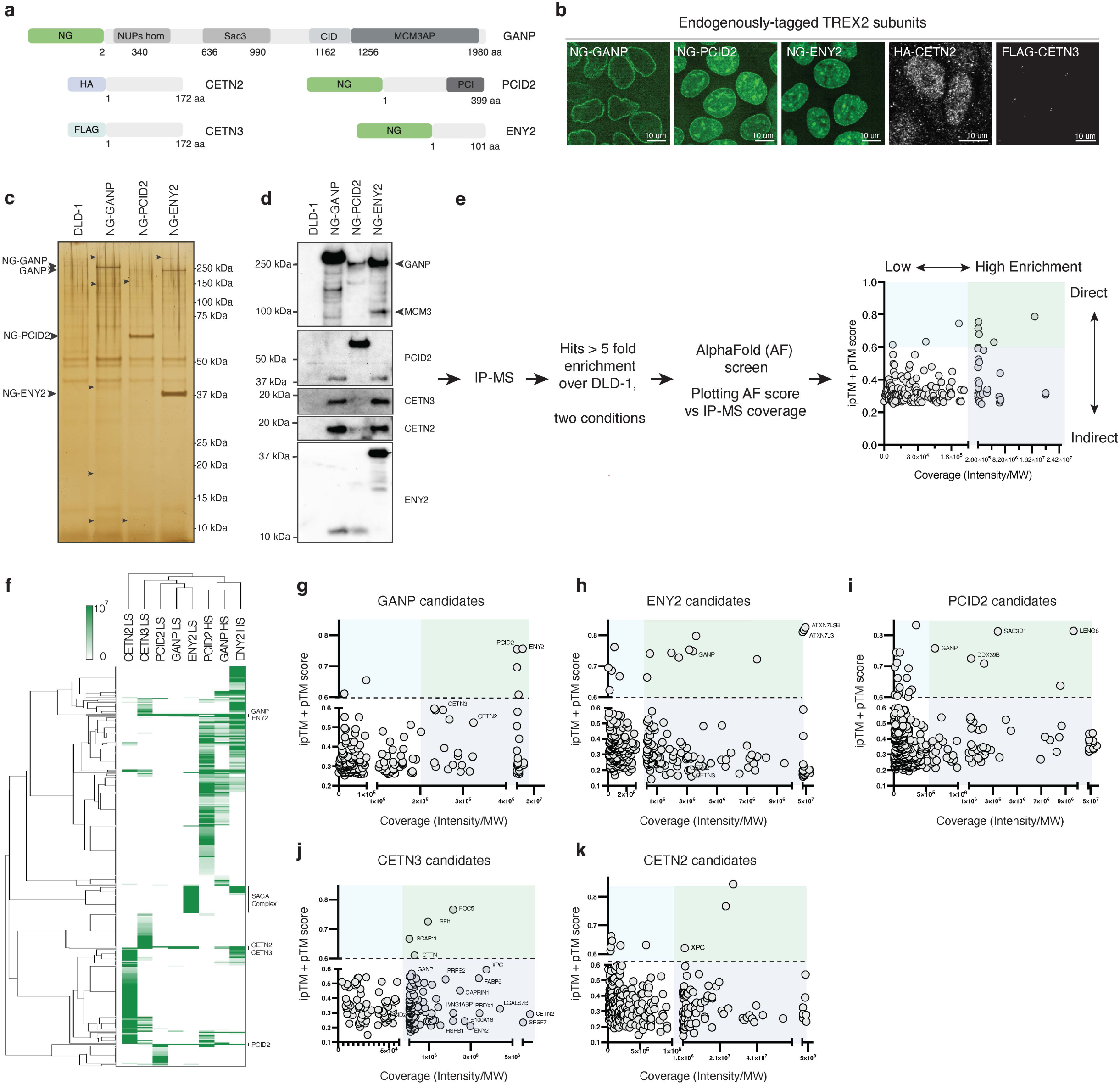
The identification of new protein partners of TREX2 subunits. **a**, A schematic of GANP, PCID2, ENY2, CETN2, and CETN3 targeting with NG, HA, and FLAG tags. **b,** Intracellular localization of NG-GANP, NG-PCID2, NG-ENY2, HA-CETN2, and FLAG-CETN3. **c**, A silver gel of NG-GANP, NG-PCID2, and NG-ENY2 immunoprecipitated using NG-beads from DLD-1 cell line. Potential protein partners are indicated with arrows. **d,** Western blotting showing formation of the TREX2 complex and corresponding levels of GANP, PCID2, CETN3, CETN2, and ENY2 binding to precipitated protein in each lane. **e**, A diagram of IP-MS-AF analysis of TREX2 protein partners. Hits with ipTM and pTM scores over 0.6 are classified as potential direct protein partners (blue sector), hits scored over 0.6 and MS coverage equivalent to TREX2 subunits as a potential direct stable interacting partner (green sector) and hits with score lower than 0.6 but high MS coverage as a stable indirect component of the complex (purple sector). **f,** A heatmap of TREX2 subunits protein partners using Hierarchical clustering and Kendall average. TREX2 subunits and SAGA complex are highlighted with the solid bar. **g-k,** Scatter plots of TREX2 subunits binding partners after IP-MS-AF screen. Representative examples of IP-MS-AF data are shown for GANP (g), ENY2 (h), PCID2 (i), CETN3 (j), and CETN2 (k). See also Figure S3.

To identify novel interacting partners, we purified each subunit using immunoprecipitation (IP) under both high and low salt extraction conditions with anti-NG, -HA or -FLAG antibodies, followed by mass spectrometry (MS) (Figure 3c-e and S3a, Table S3). Interacting partners that exhibited more than five-fold enrichment over the control were analyzed with AlphaFold Pulldown (AF) (22). In each case, the bait used for IP was utilized as bait for AF, while each interacting partner detected in the MS was utilized as prey (Figure 3e). Modelling of previously characterized complexes, such as the SAGA complex and TREX2 itself, helped us to predict the nature of interactions along two dimensions: The first dimension was binding affinity, where direct interactions had high AF (ipTM + pTM) scores (AF>0.6) versus indirect interactions with lower scores (AF<0.6). In the second dimension, abundant complexes showed high MS enrichment versus low enrichment for less abundant complexes (Figures 3e and S3b-e). This strategy, which we refer to as the quartile approach, helped us find the “best needles” in a haystack of MS data.

In total, we detected 1346 direct or abundant interactors among all TREX2 subunits. The largest number of interacting partners was observed under high-salt conditions (Figures 3f-k and S3f-h), consistent with the prediction that these conditions would more efficiently extract TREX2 proteins from the nuclear fraction. Notably, 19 partners co-purified with all TREX2 subunits (Figure S3i), including the TREX2 constituents themselves. In contrast, 18, 122, 96, 127, and 84 candidates exhibited unique specificity for GANP, PCID2, ENY2, CETN2, and CETN3, respectively. Additionally, a total of 119 candidates were found with more than one subunit but not all (Figure S3i). Thus, we identified multiple previously uncharacterized interactors of the TREX2 complex subunits. We were particularly interested in candidate proteins whose profiles suggested direct and abundant interactions that had not been previously described, either with the entire TREX2 complex or with an individual subunit. In the latter case, we speculated that such partners might reflect activities of subunits outside the context of the TREX2 complex.

### Binding partners for individual TREX2 subunits

To assess the output from the quartile plots, we first analyzed candidates that had been previously reported whose detection could serve to validate our approach (Table S4). The TREX2 subunits were used as starting points to establish the thresholds for the quartile analysis. As anticipated, the subunits that directly interact with the scaffolding subunit GANP demonstrated relatively high AF scores: 0.76 for ENY2, 0.75 for PCID2, 0.59 for CETN3, and 0.52 for CETN2. All TREX2 subunits were highly enriched in MS and were used to define the enrichment threshold (see Figure legend 3e).

Likewise, two subunits from the deubiquitinating (DUB) domain module of the SAGA complex, ATXN7L3 and USP22, have been reported to interact with ENY2 directly (23). We found both ATXN7L3 and USP22 among the ENY2 interaction candidates (Figures 3h and S3c, g). These two SAGA subunits showed AF scores of 0.80 and 0.77, respectively, and both exhibited high MS coverage, confirming their direct and abundant interactions with ENY2. By comparison, other subunits of the SAGA complex showed high MS coverage, but gave AF scores less than 0.6, suggesting that they might have high probability but indirect associations to ENY2. The cytoplasmic paralog of ATXN7L3, ATXN7L3B, has similarly been reported to associate directly to ENY2 (23). Consistent with expectations, ATXN7L3B showed an AF score of 0.83 and high MS coverage (Figures 3h and S3c). Thus, we found that the quartile approach categorized proteins previously shown to associate to individual TREX2 subunits with a high fidelity to expectations.

High affinity and high abundance candidates (green, blue and purple sectors, Figure 3e) were subjected to analysis through the STRING database, providing k-means clustered maps for each bait (Figure 4). STRING analysis suggested that each of the TREX2 subunits interacts with proteins involved in a variety of cellular processes. Interestingly, pairwise comparisons of the cellular functions between TREX2 subunits indicates overlapping but not identical pathway associations.

**Figure 4.**
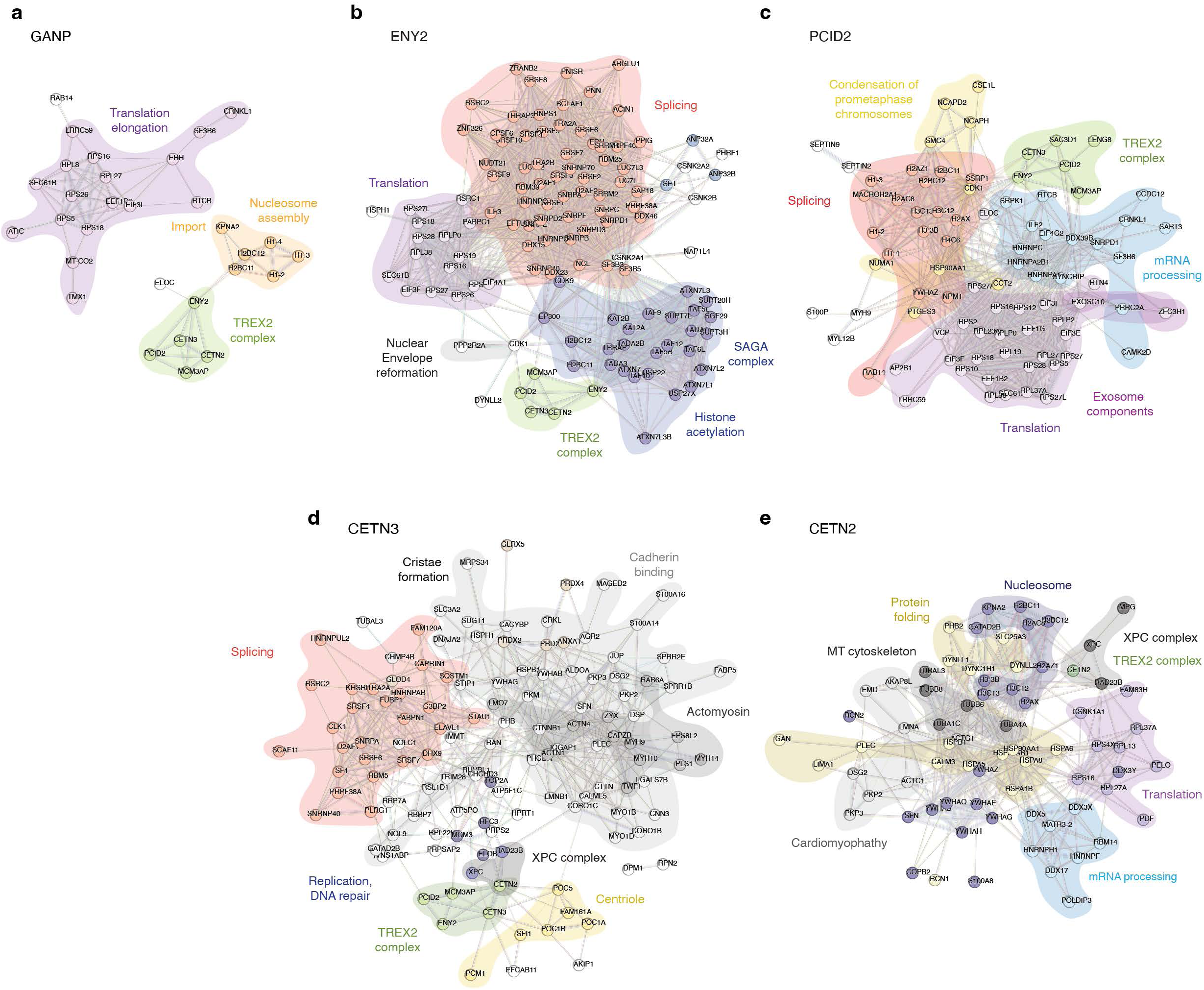
TREX2 subunits have overlapping but not identical interacting partners. STRING maps of GANP (a), ENY2 (b), PCID2 (c), CETN3 (d), or CETN2 (e) associated protein partners with AF score higher than 0.6 and >5-fold enrichment over control.

### Novel PCID2 interacting partners

We next looked for novel candidates whose interactions with TREX2 components had not been reported. We found 5, 13, 9, 8, and 2 novel and potentially direct (AF > 0.6) interacting partners of GANP, PCID2, ENY2, CETN2, and CETN3, respectively (Table S4). Our remaining analysis focused on new binding partners of the PCID2 subunit, given that it plays an essential role in bulk RNA export (Figure 1e) and that its function is poorly understood. Among the 13 novel PCID2 direct interaction candidates, database listings indicated that they function in a variety of processes, including mRNA processing, splicing, transcription elongation, RNA decay and translation initiation (Figures 3i, 4c, S3h, Table S4). Two proteins among the top PCID2 interacting partners shared intriguing sequence homology with GANP: Leukocyte Receptor Cluster Member 8 (LENG8) and SAC3 Domain Containing 1 (SAC3D1) (Figures 3i and S3h), prompting us to examine them further. Like GANP, LENG8 and SAC3D1 possess SAC3-homology domains with a PCI-fold. The LENG8 and SAC3D1 SAC3 domain share 26.7% and 52.3% similarity to human GANP, respectively (Figure 5a). Evolutionary analysis supports that the SAC3-homology domain in GANP, LENG8, and SAC3D1 have evolved from a common ancestor (Figure 5b).

**Figure 5.**
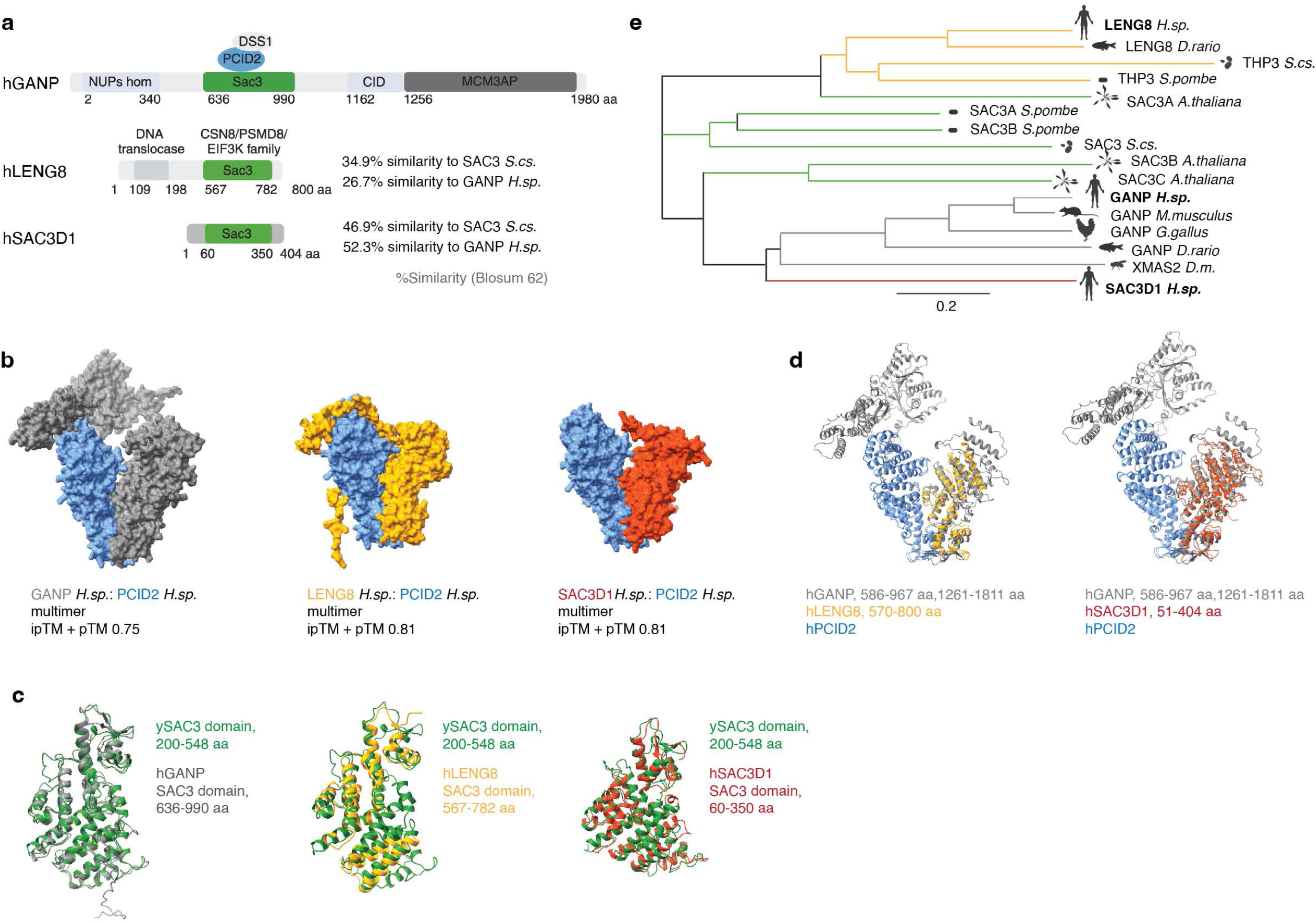
An evolutionary conserved SAC3-based subcomplexes of PCID2 subunit. **a**, A schematic of human GANP, LENG8 and SAC3D1. Similarity of proteins to *S.cs.* and *H.sp.* is shown respectively. **b**, Unrooted distance tree of GANP, SAC3D1, LENG8, SAC3, and THP3 proteins showing the evolutionary relationship (Neighbor joining) of these proteins across species. Scale bar represents number of amino acid substitutions per site. **c**, 3D alignment of SAC3 domains between human GANP, LENG8, SAC3D1, and yeast SAC3 protein. Yeast SAC3 domain is colored in green. **d**, AF multimers of human PCID2 with GANP, LENG8, or SAC3D1. **e**, 3D alignment of human GANP:PCID2 and SAC3D1:PCID2 multimers. PCID2 is colored in blue, LENG8 in yellow, SAC3D1 in red, and GANP in grey.

AF analysis of predicted binding interactions suggests that GANP, LENG8 and SAC3D1 all bind PCID2 via the PCI-fold within their SAC3 domains (Figure 5d). The overlay of predicted 3D conformations of SAC3 domains from GANP, LENG8, and SAC3D1 on the ySAC3 domain illustrates the extensive conservation of the SAC3-homology domain across these proteins (Figure 5c, e). Notably, the use of the same interface with PCID2 in each case raises the question of whether binding of GANP, LENG8 and SAC3D1 may be mutually exclusive. To address this question, we endogenously and homozygously tagged LENG8 and SAC3D1 with NG- and HA-AID in DLD-1 cells (Figures S2h-i and S5a) to analyze their interaction networks. We then assessed interacting partners of LENG8 and SAC3D1: We purified NG and HA-tagged LENG8 and SAC3D1, followed by MS and AF analysis of their interaction partners (Figures 6a-c, S5b-h, Tables S4-5). The MS datasets indicated that LENG8 and SAC3D1 interact with PCID2, as expected, but that neither of them associated with each other (Figure 6d, e). As noted earlier, neither LENG8 nor SAC3D1 were identified as GANP binding partners (Figure 3g), and neither GANP, ENY2, CETN2 nor CETN3 were found among their interaction partners (Figure S4).

**Figure 6.**
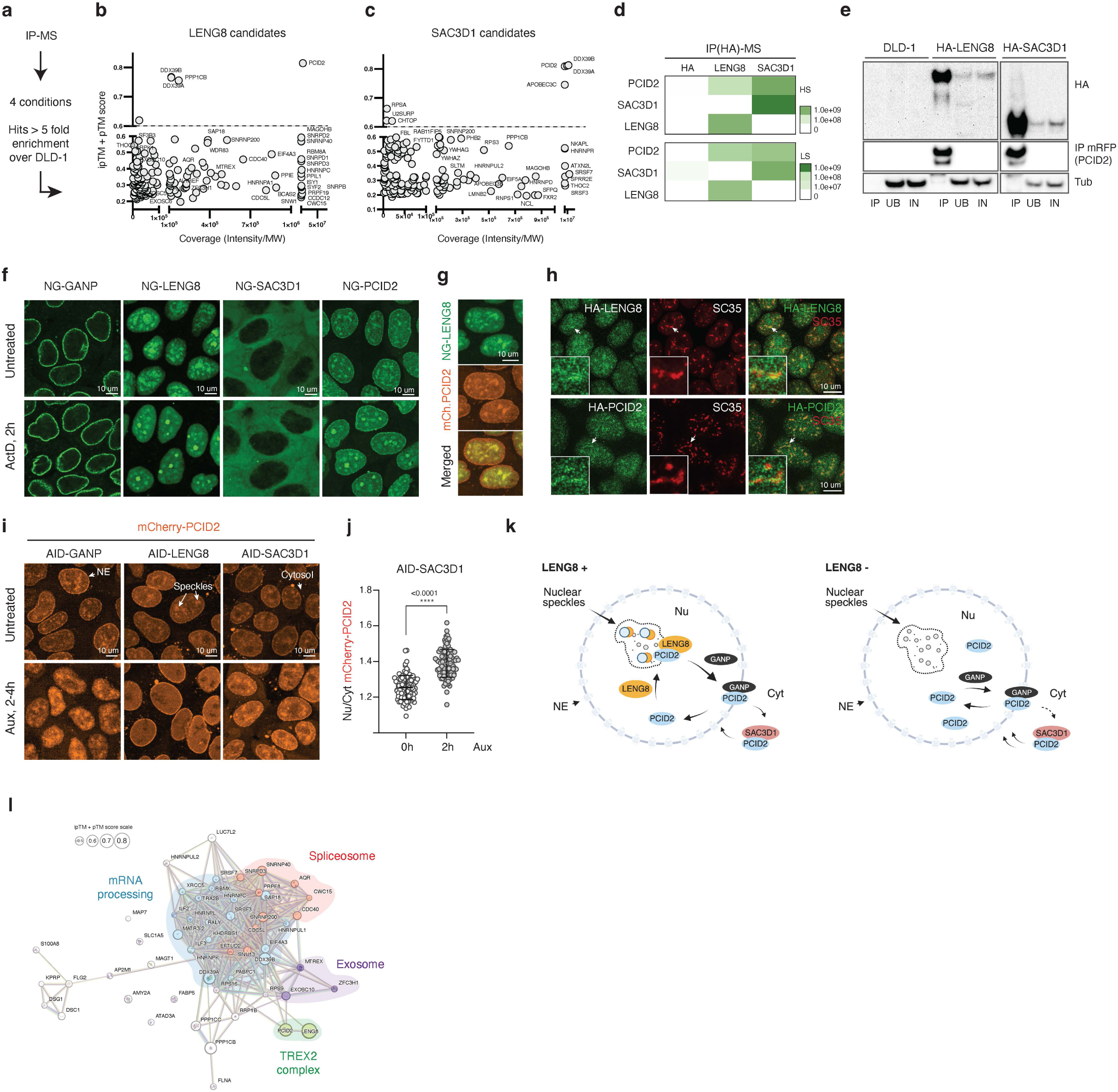
PCID2 subcomplexes with GANP, LENG8, and SAC3D1. **a-c,** A schematic of the screen and quartile plots of LENG8 (b) and SAC3D1 (c) IP-MS-AF interaction partners. **d**, Heat maps of differential abundance of PCID2, SAC3D1, and LENG8 proteins in HA-LENG8 and HA-SAC3D1 IP-MS. Green color indicates protein abundance enrichment under High Salt (HS) and Low Salt (LS) conditions. **e**, Western blot analysis of PCID2 bound HA-tagged LENG8 and SAC3D1 proteins upon immunoprecipitation. DLD-1 cell line was used as a control of RFP-beads background binding. IP – Immunoprecipitation, UB – Unbound, IN – Input, HA - Influenza hemagglutinin. **f**, Localization of NG-GANP, NG-LENG8, NG-SAC3D1, and NG-PCID2 proteins in untreated live cells and 2 h after Actinomycin D treatment. **g**, Co-localization of NG-LENG8 and mCherry-PCID2 in live cells. **h**, Co-localization of HA-tagged LENG8 and PCID2 with speckle marker SC35. **i**, Localization of mCherry-PCID2 in untreated cells and after acute loss of GANP, LENG8, or SAC3D1. Arrows indicate nuclear envelope (NE), speckles, and cytosolic localization of PCID2. **j**, Quantification of nuclear-to-cytoplasmic (Nu/Cyt) mCherry-PCID2 distribution. ****p-value < 0.0001, n = 103 untreated and n = 109 auxin-treated cells, unpaired Student’s *t*-test. Data are presented as mean values; error bars are SD. **k**, A model illustrating the regulation of PCID2 localization through its interactions with GANP, LENG8, or SAC3D1. PCID2 bound to the GANP protein localizes to the NE, but upon the loss of GANP, PCID2 is displaced into the nucleus. LENG8 localizes PCID2 within nuclear speckles, where both PCID2 and LENG8 co-localize with nuclear speckle markers, splicing regulatory factors, and DDX39B. LENG8 loss displace PCID2 from the nuclear speckles. The binding of PCID2 to SAC3D1 in the cytosol maintains the nuclear level of PCID2. **l**, STRING map of LENG8 interacting partners reproduced in at least two purification conditions with >5-fold enrichment. Circles corresponding to proteins with AF scores 0.6 and higher are enlarged respectively.

We made additional analysis of the proteins interacting with LENG8 and SAC3D1 that were reproduced in at least two independent replicates (Figures 6b-c and S5c-h) using the Human Protein Atlas (HPA) and STRING databases. We searched for protein localization of candidates and GO terms associated with their function. LENG8 bound to multiple proteins involved in splicing, mRNA processing and exosome function (Figure 6l); whereas SAC3D1 mostly interacted with mRNA processing and export proteins (Figure S7a). Interestingly, both LENG8 and SAC3D1 bound to the ATP-dependent DDX39B RNA helicase, as has previously been observed for the GANP-PCID2 complex (14). We analyzed predicted PCID2-LENG8-DDX39B and PCID2-SAC3D1-DDX39B subcomplexes using AF. AF predicted the formation of trimeric complexes with high probability in each case, with scores of 0.83 and 0.85 respectively (Figure S7f). These observations suggest that the RNA helicase DDX39B may be a shared component of complexes between PCID2 and each of the three SAC3 domain proteins.

Together, our findings indicate mutually exclusive PCID2 interactions with GANP, LENG8 and SAC3D1, suggesting that PCID2 may function outside of its role in TREX2 through alternative complexes with LENG8 and SAC3D1 plus DDX39B.

### Analysis of LENG8 and SAC3D1

LENG8 and SAC3D1 have not been well characterized, so we analyzed their behavior and function further. We examined the intracellular localization of LENG8 and SAC3D1 in comparison to GANP. Notably, GANP, LENG8, and SAC3D1 showed distinct localization (Figure 6f). GANP was predominantly localized at the nuclear envelope. In contrast, LENG8 was localized in the nucleus and in intranuclear clusters. SAC3D1 was mainly found in the cytosol, although a small fraction was observed in the nucleus (Figure S6a-c). PCID2 was abundantly found at the nuclear envelope, at intranuclear clusters and within the nucleoplasm, as well as at low abundance in the cytoplasm, consistent with the possibility that it could bind each of the three SAC3 domain proteins.

To investigate the presence of LENG8 and PCID2 in intranuclear clusters, we created a dual-tagged cell line and tracked the localization of NG-LENG8 and mCherry-PCID2. We observed that LENG8 and PCID2 co-localized inside the nucleoplasm and at the intranuclear clusters (Figure 6g). Furthermore, the localization of LENG8 and PCID2 within these clusters was actinomycin D (ActD)-dependent. After blocking transcription with ActD, both LENG8 and PCID2 re-localized from the smaller clusters to larger ones within 2 hours. In contrast, the subcellular localization of SAC3D1 and GANP remained unchanged following ActD treatment (Figure 6f, bottom panel). Halting transcription with transcription inhibitors leads to the redistribution and accumulation of splicing factors in enlarged and rounded speckles (24), dynamic organelles in the nucleus that harbor splicing factors. We wondered whether the intranuclear clusters containing LENG8 and PCID2 were nuclear speckles. To test the speckle localization of LENG8 and PCID2, we co-stained for both proteins and the speckle marker SC35. Our observations showed that both LENG8 and PCID2 colocalized with SC35, with their localization being somewhat peripheral to SC35 (Figure 6h). This data indicated that both LENG8 and PCID2 were targeted to the nuclear speckle compartment.

To assay how GANP, LENG8, and SAC3D1 might regulate PCID2 localization, we developed CRISPR/Cas9 AID-tagged cell lines for GANP, LENG8 and SAC3D1 with mCherry-tagged PCID2 (Figures 6i and S6d-e, (5)). We observed that GANP depletion caused loss of PCID2 from the nuclear envelope, while LENG8 depletion caused PCID2 to re-localize from intranuclear clusters to the nucleoplasm without affecting its nuclear envelope localization. In contrast, SAC3D1 depletion slightly but significantly increased the nuclear-to-cytoplasmic ratio of PCID2 distribution (Figures 6i-j and S6f). These findings suggest that PCID2 localizes at the nuclear envelope with GANP, where they may coordinately function in mRNA nuclear export. Separately, PCID2 localizes at nuclear speckles in a LENG8-dependent manner, and it appears that SAC3D1 may modulate the overall level of PCID2 in the nucleus. Thus, GANP, LENG8, and SAC3D1 have distinct impacts on PCID2’s intracellular localization (Figure 6k), further suggesting novel and previously undescribed roles for LENG8 and SAC3D1 as alternative functional partners distinct from GANP.

### LENG8 associates with spliceosome components

For the remainder of this report, we will focus on the findings regarding LENG8, since we were particularly intrigued by its possible role within nuclear speckles and since we found multiple spliceosome components among its interaction partners (Figure 6l). Notably, a total of 318 proteins were listed as spliceosome components and spliceosome-associated proteins within the HPA or as curated splicing regulators in the IRA database (25); we found that 99 of these proteins were also identified among the LENG8 interacting components (Figure S7b). Among these LENG8 and spliceosome associated partners, 31 had been indicated to localize within nuclear speckles (Figure S7c), and we confirmed the nuclear and speckle localization of 5 LENG8-bound proteins (Figure S7d).

Moreover, LENG8 interacts with multiple components of the core spliceosome machinery, which are crucial for the assembly and activation of the spliceosome (Figure S7g). Together, these results indicate that LENG8 is a speckle component that associates strongly with the spliceosome. Given that LENG8 loss impacts PCID2 recruitment to nuclear speckles, we evaluated the integrity of speckles using PNN, a protein abundantly concentrated in nuclear speckles (26). In contrast to PCID2, the localization of PNN remained visually unchanged after the loss of LENG8 (Figure S7e), suggesting that PCID2 re-localization does not reflect speckle disassembly after LENG8 depletion.

### LENG8 loss shifts distal-to-proximal polyadenylation sites usage and alternative mRNA processing

Given LENG8’s interaction with components of the spliceosome machinery, we tested whether it might play a role in RNA splicing. Notably, the previously identified LENG8 homologue in budding yeast, Thp3, forms a complex with the Csn12 and Sem1 proteins that physically interacts with core components of the spliceosome, U1-U2 snRNPs and regulates mRNA splicing (27). We conducted bulk RNA sequencing analysis on samples depleted of LENG8, comparing them to samples depleted of SAC3D1 and PCID2 (Figures 7a and Table S2B). The depletion of LENG8 resulted in the upregulation of a small group of RNAs, primarily protein-coding genes, along with the second, smaller group consisting of long intergenic non-coding RNAs (lincRNAs) and antisense RNAs (Figure S8a). Loss of SAC3D1 significantly affected the abundance of only two transcripts. Notably, the transcriptomic profiles of LENG8 and PCID2 did not strongly overlap. The transcripts that were upregulated in the absence of LENG8 were mostly downregulated when PCID2 was absent (Figure 7a). Most of the RNA transcripts affected by the loss of LENG8 encode proteins that are involved in regulating RNA metabolic processes and DNA-templated transcription (Figure S8b-c). This indicates that, in the absence of LENG8, cells attempt to upregulate nucleic acid metabolic processes. Our findings led to a coherent view that the LENG8-PCID2 subcomplex functions independently of PCID2’s role within the TREX2 complex.

**Figure 7.**
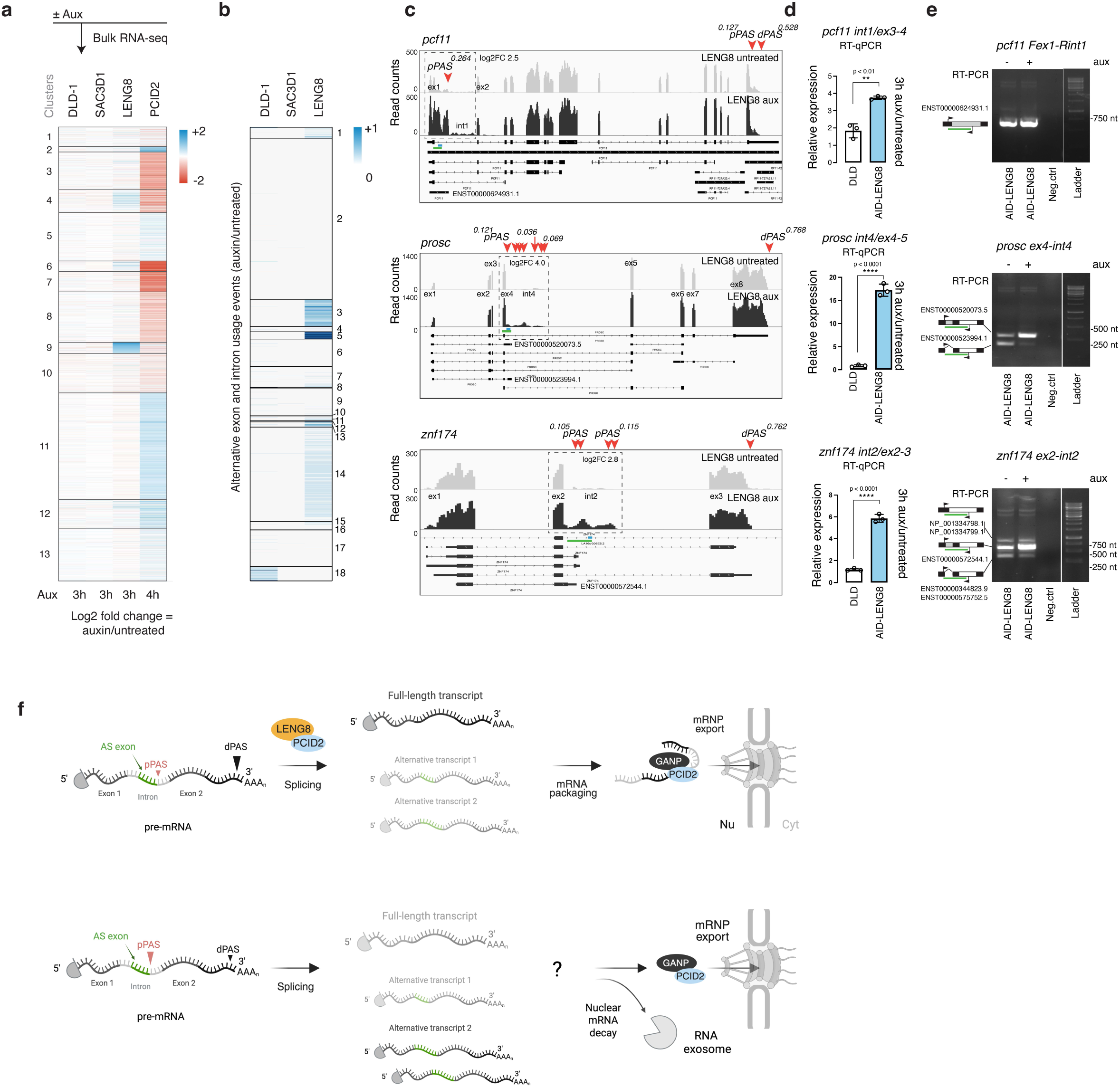
LENG8 loss alter PAS usage and mRNA processing. **a,** A heat map of unsupervised k-means clustering of differentially expressed genes 3-4 h after 5Ph-IAA or auxin treatment of cells expressing AID-tagged SAC3D1, LENG8, or PCID2. DLD-1 — parental cells treated with 5-Ph-IAA. SAC3D1 and LENG8 5-Ph-IAA-treated cells were compared to DMSO-treated cells, PCID2 auxin-treated cells were compared to untreated cells. Three independent biological replicates were used to perform RNA-seq of the indicated cell lines. **b**, A heat map of differential exon (DEU) and intron usage (DIU) events in DLD-1, SAC3D1 or LENG8-depleted cells. All alternative usage events were scaled from 0 to 1 and normalized to gene length. Events with a higher count of DEUs and DIUs will be displayed in a darker color. **c**, IGV snapshots of RNA-Seq data for *pcf11*, *prosc*, and *znf174* transcripts. Regions with alternative usage events and increased read counts are marked with dashed squares. Numbers represent the frequency of PAS usage and fold change of upregulated region. pPAS – proximal polyadenylation site, dPAS – distal polyadenylation site (dPAS). **d**, RT-qPCR analysis of upregulated regions (marked in blue line on **c**) in *pcf11*, *prosc*, and *znf174* transcripts after LENG8 loss. Graphs show mean values of three technical replicates of one experiment; error bars are SD. Asterisks indicate ****p-value < 0.0001 and **< 0.01 in unpaired two-tailed Student’s *t*-test. **e**, Agarose gel electrophoresis of the RT-qPCR products for *pcf11*, *prosc*, and *znf174* transcripts, covering the last exon before the upregulated intron and the upregulated intron itself (marked in a green line on **c**). **f**, The model of the distal-to-proximal polyadenylation sites (PAS) switch and alternative mRNA splicing. In cells that have LENG8, the distal PAS is used to generate the dominant full-length mRNA transcript. If an alternative proximal PAS exists within an intron, cells use this site to produce shorter, less abundant alternative mRNA isoform that might serve regulatory role. In the absence of LENG8, the dPAS is still utilized to create a full-length transcript. However, there is a noticeable shift towards increased usage of the pPAS. This shift leads to a higher abundance of shorter isoforms, particularly those that retain additional sequences. Fully processed transcript potentially utilizes GANP-PCID2 complex (as a part of TREX2) for the following nuclear export, whereas short isoforms might get exported from the nucleus or send to nuclear mRNA decay.

To assess whether loss of LENG8 impacts splicing, we examined differential exon usage (DEU) and differential intron usage (DIU) events in the absence of LENG8. Our analysis of alternative splicing events revealed that the number of DEU and DIU was significantly higher without LENG8 compared to those observed in the absence of SAC3D1, or in DLD-1 cell line treated with auxin (Figure 7b). A manual assessment of read counts revealed that most of these events occurred within the sequences of canonical introns and were associated with exons of alternative, often shorter isoforms that utilize a different poly(A) tail (Figures 7c and S9). In-depth examination of RNA-seq data confirmed the increased abundance of reads occurred in regions corresponding to short alternative isoforms. This increase did not impact the overall abundance of the primary longer transcript. Furthermore, we confirmed the upregulation of intronic reads that corresponded to an alternative short isoform by qPCR (Figure 7d) and confirmed the splicing junction between these intronic reads and the adjacent upstream exon using RT-PCR analysis (Figure 7e). Transcripts which demonstrated DEU and DIU events had a preference to use a proximal polyadenylation site (pPAS) instead of a distal polyadenylation site (dPAS) for cleavage and polyadenylation. We found examples of tandem (rps28), alternative exon (prosc, znf174, ing3, siva1, znf789), or intronic (pcf11) alternative polyadenylation usage events (Figures 7c and S9). This suggests a potential role for LENG8 in mRNA splicing coupled to alternative polyadenylation (Fig. 7f).

Together, these data indicate that LENG8 may be a functional homologue of Thp3, contributing to splice-site selection and the choice of alternative polyadenylation sites in the upstream regions of selected mRNAs, ultimately leading to the production of RNA isoforms with different 3’-terminal exons and UTRs.

## DISCUSSION

The subunits of the TREX2 complex do not behave identically, nor do mutations in these proteins cause equivalent phenotypes (6, 7, 16–21). To understand the different roles of individual TREX2 subunits, we tagged them with auxin-induced degrons to allow for their rapid and selective depletion. While it had been generally anticipated that loss of any TREX2 subunit should cause nuclear mRNA retention (3, 6), we interestingly found that poly(A) mRNA retention was observed for some subunits (GANP, PCID2) but not for others (ENY2 or CETNs) (Figure 1). Moreover, different patterns of transcriptomic changes after the loss of individual subunits (Figure 2) supported the idea that they do not strictly function in tandem. To assess their cellular roles in more detail, we developed an integrated pipeline that combines CRISPR/Cas9 endogenous tagging, Immunoprecipitation (IP), Mass Spectrometry (MS), AlphaFold-based modelling (AF), and bioinformatic analysis of interacting partners (Figures 3 and 4). The interaction profiles of each TREX2 subunit showed binding partners shared among all subunits, as well as partners that were unique to each subunit. As an example of such alternative complexes, we analyzed PCID2 binding to SAC3D1 and LENG8; we found that these proteins assemble functional complexes with PCID2 in a manner that is mutually exclusive to its assembly into the classical TREX2 complex. Taken together, these findings suggest that TREX2 proteins participate in a network of interactions beyond their suggested roles in nuclear mRNA export (3, 6), raising the exciting possibility that they may participate in crosstalk between many nuclear pathways that regulate gene expression.

There was a surprising level of divergence among the behaviors of individual TREX2 subunits. Beyond the distinct impacts of subunit depletion on transcriptomic patterns and mRNA export (Figures 1 and 2), endogenously tagged proteins showed distinct localizations (Figure 3b). Notably, CETN2 and CETN3 did not co-localize closely with other subunits or associate to the nuclear envelope. Co-localization among the other TREX2 subunits was stronger, with GANP localized to the nuclear rim, while ENY2 and PCID2 localized to the nuclear rim, in the nucleoplasm, and in nuclear speckles (Figure 3b). The presence of ENY2 at sites with low GANP abundance may indicate its involvement in GANP-independent functions, including its well-known role within the SAGA complex (28). Similarly, our findings suggest that PCID2’s localization at sites without GANP involves alternative complexes, such as those with SAC3D1 and LENG8 (Figure 3), which we further characterized in this report.

We found SAC3D1 and LENG8 as abundant and direct binding partners of PCID2 using a quartile approach to categorize interaction candidates in two dimensions — according to MS coverage and to AF predictions of their binding affinity (Figure 3). MS analysis further showed that GANP, SAC3D1 and LENG8 each interact strongly with PCID2 but in a manner that is mutually exclusive (Figure 5 and 6). SAC3D1 was cytoplasmic while LENG8 localized within the nucleoplasm and nuclear speckles (Figure 6f, h). It has previously been reported that PCID2 localization at the nuclear envelope requires GANP (5, 6). We observed that depletion of either SAC3D1 or LENG8 altered PCID2 accumulation at other sites (Figure 6i-j). SAC3D1 depletion shifted the balance of PCID2 from the cytoplasm to the nucleus. In *Drosophila melanogaster*, PCID2 shuttles between the nucleus and the cytoplasm; a cytoplasmic chaperone protein, NudC, promotes PCID2’s cytoplasmic stability and its activity in cytoplasmic mRNA trafficking (29). It is interesting to speculate that SAC3D1 may act like NudC and control PCID2 within the cytoplasm (Figure 6i-k), although it has not been established that SAC3D1 has chaperone activity or that human PCID2 mediates cytoplasmic mRNA transport. By contrast, LENG8 depletion released PCID2 from nuclear speckles. LENG8 did not appear to be a structural component of speckles since other speckle-associated proteins were not dispersed upon its depletion. Together, these findings suggest that PCID2’s localization within cells is modulated by its allocation between complexes with these three SAC3-homology domain proteins.

Consistent with the established role of the TREX2 complex (2, 3), depletion of either PCID2 or GANP caused poly(A) RNA accumulation within the nucleus (Figure 1), indicating a role for PCID2-GANP complexes in licensing a subset of RNA transcripts for export. The close alignment of transcriptomic changes upon depletion of PCID2, GANP or the key nuclear pore protein TPR (Figure 2a), further suggests their functional alignment in RNA export at the nuclear pore. Depletion of ENY2 or CETNs did not promote bulk poly(A) RNA retention (Figure 1), suggesting that they may not be essential at the same late stages of mRNA packaging and nuclear export.

As we looked for the function of LENG8, we noted that budding yeast PCID2-like proteins Thp1 and Csn12 form two structurally related complexes: Sac3-Thp1-Sem1 and Thp3-Csn12-Sem1, where Sac3 and Thp3 are the paralogues of GANP and LENG8 respectively. The Thp3-Csn12-Sem1 complex promotes pre-mRNA splicing (15, 27) and transcription elongation (30). Csn12 interacts with core spliceosome components (31) and plays a functional role in mRNA splicing (27). By contrast, the Sac3-Thp1-Sem1 complex has higher affinity to mRNA and facilitates mRNA nuclear export (15). We speculated that the PCID2-LENG8 and PCID2-GANP interactions might fulfill the roles of Thp3-Csn12-Sem1 and Sac3-Thp1-Sem1, respectively. The Sac3-Thp1-Sem1 (13), Thp3-Csn12-Sem1 (15), and GANP-PCID2-DSS1 (32) complexes exhibit an inverted V-shape structure, where juxtaposition of winged helix (WH) domains generates a positively charged surface that play a key role in nucleic acids binding (15). Both PCID2-LENG8 and PCID2-GANP complexes exhibit a similar V-shape structure (Figure 5d, (12, 14, 32)), with the positively charged surface that may, similarly to Thp3-Csn12 and Sac3-Thp1, provide a platform for interaction with RNA. The evolutionary relationship between Thp3 and human LENG8 (Figure 5b), together with LENG8 localization in nuclear speckles and its interactions with spliceosomal and mRNA processing factors (Figure 6l), further supported the idea that LENG8 might participate in a complex with functions similar to those of Thp3-Csn12-Sem1.

Consistent with the idea that human LENG8-PCID2 complexes modulate RNA processing, we found changes in splice site selection and polyadenylation of transcripts upon its depletion (Figure 7c). Interestingly, one mRNA whose processing is markedly changed after LENG8 depletion is PCF11, encoding a low abundance cleavage and polyadenylation factor known for its role in transcription termination and early polyadenylation (33). Modulation of PCF11 expression after LENG8 depletion could trigger further alternative mRNA processing, including genome-wide polyadenylation site usage. Moreover, we and others found DDX39B as a component of the LENG8-PCID2, GANP-PCID2, and SAC3D1-PCID2 subcomplexes (Figures 3i, and 6b, c, (12, 14)). GANP-PCID2 and LENG8-PCID2 complexes are believed to function as disassembly factors for DDX39B-containing mRNPs (12). Considering that DDX39B is required for the release of spliced transcripts from nuclear speckles (34) and our findings that PCID2, LENG8, and DDX39B are all localized in nuclear speckles, it appears plausible that LENG8-PCID2-DDX39B activity may promote mRNA remodeling within the speckle domain. These findings do not exclude a role of LENG8-PCID2 in nuclear RNA decay processes (35, 36), and indeed we found components of the exosome machinery, including MTREX, EXOSC10, and ZFC3H1, as potential direct interacting partners of PCID2 and LENG8 in our IP-MS-AF analysis (Figure 6l). Our data may thus suggest that the LENG8-PCID2 complex is part of an intricate autoregulatory circuit interlinking mRNA processing, decay and export.

In conclusion, we find that TREX2 subunits form protein complexes beyond TREX2 itself, and that such interactions link multiple aspects of nuclear mRNA modification and export. It will be of considerable interest to investigate how TREX2 subunits are exchanged between these interaction partners and how these exchanges might facilitate correct RNA maturation and delivery to the cytosol.

### Limitations of the study

We performed IP-MS using the colorectal adenocarcinoma cell line DLD-1. It is important to note that these cells may not fully represent the surrounding protein environment of TREX2 subunits, LENG8, and SAC3D1 in other tissues, nor do they account for the dynamic changes in the proteome that occur during development. Additionally, we utilized only two variants of extraction conditions for immunoprecipitation, which may not have effectively purified all proteins.

The introduction of an AID degron and the integration of TIR1 could lower overall protein levels, potentially impacting the manifestation of phenotypes upon loss of AID-tagged proteins. In addition, we only tagged one terminus of the protein; therefore, any additional short isoforms located on the opposite end will not be visualized or degraded. Despite this limitation, the localization of our NG or AID-tagged cell lines was consistent with previously published data, and we confirmed that AID-proteins were fully degraded using endogenous antibodies. We confirmed the splicing junction between upregulated intronic reads and the adjacent upstream exon; PAS sequencing or 3’ RACE is required to determine the whole sequence of the short isoforms produced upon LENG8 loss.

## Resource availability

### Lead contact

Requests for further information, resources, and reagents should be directed to the lead contacts, Vasilisa Aksenova and Mary Dasso.

### Materials availability

All materials will be available upon reasonable request.

### Data and code availability

The mass spectrometry proteomics data have been deposited to the ProteomeXchange Consortium via the PRIDE (37) partner repository. These data sets are currently private. The sequencing data have been deposited in NCBI Gene Expression Omnibus (GEO). This data set is currently private. The data underlying the main and supplementary figures will be provided as a Source Data file. The authors will provide all other data upon reasonable request. The authors utilized standard, published, open-source tools for processing RNA-seq, MS, and AF data. The authors acknowledge the use of OpenAI’s ChatGPT (GPT-5) for assistance in writing and refining minor data analysis scripts, which will be included in the Source Data file.

## Acknowledgments

VA, AA, EG, and MD were supported by the Intramural Research Program of the Eunice Kennedy Shriver National Institute of Child Health and Human Development at the National Institutes of Health, USA (Intramural Project #Z01 HD008954). We are grateful to Tianwei Li, James Iben (NICHD Molecular Genomics Core) and CE and Ryan Dale (Bioinformatics and Scientific Programming Core) for help with the library construction, paired-end sequencing and bioinformatics support. This work utilized the computational resources of the NIH HPC Biowulf cluster. We also thank Wolfgang Resch for his help in setting up the AlphaFold analysis on the Biowulf cluster. Lastly, we appreciate the members of Mary Dasso’s group for their valuable reviews and constructive feedback on the manuscript.

## Author contributions

VA and MD developed hypothesis and designed experiments. All experiments, analyses, and project development were conducted by VA, except for MS, which was done together with AA. FISH was done together with EG. EG was also responsible for generating the DLD-1 CETNs cell lines. VA supervised EG throughout the project. CE performed the bioinformatics analysis of the RNA-seq data. VA and MD wrote the paper. AA, EG, CE edited manuscript.

## Methods

### Gene targeting

The CRISPR/Cas9 system was used for endogenous targeting of all genes at their N-termini. TPR, GANP, and PCID2 have been targeted as previously described (5, 38, 39). ENY2, CETN2, CETN3, LENG8, and SAC3D1 were tagged either with a minimal functional AID (1xmicroAID) 71-114 amino-acid (40) and HA tag; or 1xmicroAID and FLAG (CETN3); or NeonGreen (NG) fluorescent protein. The DNA sequences of 1xmicroAID, HA, FLAG, and NG tags were codon-optimized and synthesized (IDT or Genewiz). The homology arm sequences were amplified from genomic DNA extracted from DLD-1 cells and integrated into a universal donor vector modified from one previously described (5). cDNAs of mCherry, puromycin, hygromycin, and TIR1 sequence were amplified as previously described (5). The RCC1 (NC_000001.11, Regulator of Chromosome Condensation 1) locus was chosen to knock-in TIR1. In brief, the RCC1 locus was endogenously targeted with a construct containing an infra-red fluorescent protein (IFP2.0), 9Myc-TIR1, and a blasticidin resistance gene sequence, as described in (5). TPR, GANP, PCID2, ENY2, CETN2, and CETN3 AID-tagged cell lines were tagged with OsTIR1 amplified from pBABE TIR1-9Myc (Addgene #47328)(41); whereas LENG8 and SAC3D1 were tagged with OsTIR1_74G amplified from pMK381 (Addgene, #140536)(42). The gRNAs for ENY2, CETN2, CETN3, LENG8, and SAC3D1 were designed using https://crispor.gi.ucsc.edu/ and integrated into pX330 (Addgene #42230) vector using Zhang Lab General Cloning Protocol (43). All PCR reactions were performed using Platinum SuperFi (ThermoFisher, 12351010) or Hi-Fi Taq (ThermoFisher, 11304011) DNA polymerases. All fragments for genetic cloning were purified using Gel and PCR Clean-up kit (Takara Bio, 740609.250). Oligonucleotide sequences of gRNA and primers for amplification of homology arms are listed in Supplementary Table 1. Additional details are provided in Supplementary Fig. 2.

### Genotyping

Tagged clones of ENY2, CETN2, CETN3, dual CETN2/3, LENG8, and SAC3D1 cell lines were genotyped using two primer sets. The first set of primers (F*in* and R*in*) was used for initial screening of the clones. The second set of primers (F*out* and R*out*), listed in Supplementary Table 1, was used to confirm homozygosity of the selected clones. A schematic illustrating primer annealing is shown in Supplementary Fig. 2a.

### Cell culture

The human colorectal cancer cell line DLD-1 was cultured in DMEM media (ThermoFisher, 10313-021) supplemented with heat-inactivated 10% FBS (R&D system Biotechne, S11550), 2 mM GlutaMAX (ThermoFisher, 35050-061), 100 IU/ml penicillin and 100 μg/ml streptomycin (ThermoFisher, 15140-122), and in 5% CO_2_ atmosphere at 37°C.

### Plasmid DNA preparation and transfection

Transfection of DLD-1 cell line was performed as previously described (5, 38). Briefly, for transfection 0.1 x 10^6^ cells/well were plated in 12-well plates a day before transfection. Plasmids for transfection were prepared using the NucleoSpin kit (Takara Bio, 740588.20). DLD-1 cells were transfected with 500 ng of donor and 500 ng of mixture of gRNA plasmids at a ratio 1:1 using ViaFect transfection reagent (Promega, E4981) according to the manufacturer’s instruction. Seventy-two hours after transfection, cells were seeded on 10-cm dishes with the selective antibiotics hygromycin 200 μg/ml (ThermoFisher, 10687010), or blasticidin 10 μg/ml (ThermoFisher, A1113903), or puromycin 3 μg/ml (ThermoFisher, A11138-03) until clones were formed on a plate. Clones were genotyped using primer sets in Supplementary Table 1. Homozygous endogenously tagged clones were subsequently used to analyze the localization and expression of HA-, FLAG-, NG-, or mCherry-targeted proteins. Healthy growing clones with proper molecular weight and localization of tagged protein were propagated further for downstream analysis in regular complete media without selective antibiotics.

### Immunoprecipitation (IP) and Mass Spectrometry (MS)

Two alternative methods were used for TREX2 subunits extraction for IP and MS.

### High salt extraction

Unsynchronized DLD-1 cells were grown on tissue culture-treated 150×20 mm dishes for 2 days to reach 80% confluency before protein purification. To extract nuclear-and nuclear envelope enriched proteins, we used High Salt (HS) buffer, HSA (10 mM TrisHCl, pH7.5; 0.5M NaCl, 0.3% Triton X100, 5 mM MgCl_2_, 2 mM DTT, containing EDTA-free protease inhibitor cocktail (Roche, 04693159001). In brief, cells quickly washed with PBS and scraped directly from plates in 1 ml of buffer HSA per dish, 4 dishes per cell line were used for single IP experiment. Cell lysates were homogenized 10 times by syringe with a 21-gauge needle, collected in low retention 1.5 ml tubes (Fisher Scientific, 02-681-320) and sonicated for 15 cycles (30 sec ON/ 30 sec OFF) at 4^0^C. Samples were transferred to centrifuge tubes (Beckma Coulter, 343776), ultra-centrifuged at 500 000 (TLA-120.1 rotor) for 10 min at 16^0^C and supernatants were collected back to low retention tubes. To return to physiological ionic strength (150 mM NaCl) samples were diluted with buffer HSB (10 mM TrisHCl, pH7.5; 5 mM MgCl_2_, 2 mM DTT, containing EDTA-free protease inhibitor cocktail (Roche, 04693159001). Diluted samples were incubated on ice for 60 min. NG-Trap Agarose (Chromotek), RFP-Trap Agarose (Chromotek), anti-HA magnetic (Pierce, 88836), and anti-FLAG M2 magnetic (Sigma, M8823) beads were used to immunoprecipitate NG-, mCherry-, HA-, or FLAG- endogenously tagged proteins, respectively, at a ratio of 20 µl of beads per dish. Beads were washed 4 times with buffer HSC (10 mM TrisHCl, pH7.5; 0.15M NaCl, 0.05% Triton X100, 5 mM MgCl_2_, 1 mM DTT) and blocked with 2% gelatin made on HSC buffer before proceeding to overnight IP at 4^0^C. The following day, beads were washed 4 times to remove nonspecific binding with HSD buffer (10 mM TrisHCl, pH7.5; 0.15M NaCl, 0.1% Triton X100, 5 mM MgCl_2_, 2 mM DTT) on Poly-Prep Chromatography Columns (BioRad, 731-1550). Samples in the last washed were moved to new tubes and boiled for 15 min at 98^0^C along with input (IN) and flowthrough samples (UB). From IP, 10% of beads was saved for silver staining, and the rest of the beads were used for IP-WB. Equal amounts of IN and UB samples were taken from the lysates and loaded onto SDS–PAGE. To enable efficient separation and transfer of high–molecular-weight proteins we used 10-well NuPAGE 4-12% Bis-Tris gels (ThermoFisher, NP0321BOX), NuPAGE MOPS SDS running buffer (ThermoFisher, NP0001-02), Immunobilon-B PVDF 0.45 um membrane (Millipore, IPVH304F0), and 10% MeOH-containing NuPAGE transfer buffer (ThermoFisher, NP0006-1); for low–molecular-weight proteins we used 10-well 8% Bolt Bis-Tris Plus Wedge Well gels (ThermoFisher, NW00080BOX), Bolt MES SDS running buffer (ThermoFisher, B0002-02), 0.1 um NC nitrocellulose blotting membrane (Amersham Protran, 10600000), and 20% MeOH- containing NuPAGE transfer buffer (ThermoFisher, NP0006-1), using semidry transfer (TE77XP SemiDryBlotter, Hoefer). Efficacy of immunoprecipitation was analyzed with antigen-specific antibodies.

### Low salt extraction

To prepare cell protein lysate with Low Salt (LS) buffer or LSA, cells were scraped in media, transferred to 15 ml tubes, pelleted at 300g for 5 min, washed 1 time with PBS, and pelleted again at 300g for 5 min. Cell pellets from one 150×20 mm dish were resuspended in 1 ml of buffer LSA (20mM HEPES, 0.5% NP-40, 100 mM KCl, 1 mM MgCl_2_), containing 200 nM okadaic acid, 10 µg/ml DNAseI, 10 µg/ml each leupeptin (Sigma, L8511), pepstatin (Sigma, P4265) and chymostatin (Sigma, C7268), and 1x RNAsecure RNAse inhibitor (ThermoFisher, AM7006). Lysates were incubated at 4^0^C for 10 min, homogenized 10 times by syringe with a 21-gauge needle, collected in 1.5 ml tubes (Fisher Scientific, 02-681-320) and sonicated for 15 cycles (30 sec ON/ 30 sec OFF) at 4^0^C. Lysates were spun down for 15 min at 19000g at 4^0^C in bucket Eppendorf rotor. Supernatants were incubated with corresponding beads for 3h at 4^0^C on a rotator. Beads were washed 3 times in 1 ml of buffer LSA, pelleted each time at 400 g for 10 sec in a swinging bucket eppendorf rotor (or on magnet separator), transferred to a new tube and washed 2 times again as above. From IP, 10% of beads were used for silver staining. Silver staining was performed using Pierce Silver Stain for Mass Spectrometry (ThermoFisher, 24600). All other steps for HLA and LSA samples were identical and matched the description provided above.

Samples were individually analyzed with LC-MS for label-free quantification according to the manufacturer’s instructions. The mass spectrometric analysis was conducted on an LTQ Orbitrap Lumos (ThermoFisher Scientific) based nanoLCMS system. Protein identification and quantitation analysis were carried out on the Proteome Discoverer 2.2 platform (ThermoFisher Scientific). Peptide IDs were assigned by searching the resulting LC-MS raw data against UniProt/SwissProt Human database using the Mascot algorithm (V2.6, Matrix Science Inc.). Peptide-spectrum matches (PSM) were further validated with the Percolate algorithm. We determined the protein-level intensity based on the median of peptide-level intensity from the Proteome Discoverer-produced abundances. Intensity of relevant proteins in IP of TREX2 subunits and LENG8 and SAC3D1 are shown in Supplementary Table 3 and Supplementary Table 5, respectively.

### Westen Blotting (WB) of total cell lysates

For total cell lysates DLD-1 cells were grown on 6-well plates and pellets were lysed in 2x Laemmli Sample Buffer (Bio-Rad, #1610737), boiled for 15 min at 98^0^C, and ultra-centrifuged at 500 000 (TLA-120.1 rotor) for 10 min at 16^0^C. SDS-PAGE and Western Blotting were performed as described above. Total protein abundance was assessed using Coomassie Blue staining of SDS-PAGE to normalize sample loading. The PVDF and nitrocellulose membranes were blocked in 5% non-fat milk (Carnation Nestle) or 2% hydrolysate gelatin enzymatic (Sigma, G0262), respectively, for 1 h, followed by overnight incubation with the primary antibody at 4^0^C. The membrane was rinsed and probed for 2 h at RT with the secondary antibodies conjugated to HRP (Thermofisher, anti-rabbit A16104, anti-rat 31470, or Sigma, anti-mouse NA931V) with a dilution of 1:15000 in 1x TN buffer (150 mM NaCl, 10 mM TrisHCl, pH 7.5, 0.1% Tween 20). The signal from antibodies was visualized with ECL Western Blotting Substrate (Promega, W1015) or SuperSignal West Pico PLUS Chemiluminescent substrate (ThermoFisher, 34578) using FluorChem Imaging System (ProteinSimple).

### AlphaFold analysis

All computational work was performed on NIH HPC Biowulf cluster (https://hpc.nih.gov). The AlphaPulldown (AP) package (22) was used for the first round of protein-protein interactions prediction. We generated 10 models for each multimer and assessed their accuracy by calculating the sum of the interface predicted template modeling (ipTM) scores and the predicted template modeling (pTM) scores. These ipTM and pTM sums were then plotted against the protein coverage observed in mass spectrometry (MS). All multimers with a score higher than 0.6 were re-analyzed using AlphaFold version 2.3.2(44). Scores below 0.6 in Supplementary Tables 3 and 5 were generated using the AP method, while scores above 0.6 represent an average of the AP and AF scores.

The confidence of the predictions was validated using a control set of interacting partners from the SAGA complex (Supplementary Fig. 3c-e). SUPT20H, TAF5L, and SUPT3H subunits served as bait and subunits from the Core, HAT, SPL, TRRAP, and DUB domains were used as prey. Cryo-EM maps and contact residues of the interacting surfaces of the SAGA complex were obtained from the 7KTS and 8H7G PDB files and visualized in Chimera version 1.7. We found that when contact residues were visualized in Chimera, this consistently corresponded to an AF score between 0.7 and 0.9 and indicated a substantial interaction interface between the two proteins. When the interacting surface involved smaller protein regions, AF scored this interaction between 0.3 and 0.7. Non-interacting proteins showed no contact residues in Chimera. Representative examples with large, medium, or minimal interacting surfaces among SAGA complex subunits are shown in Supplementary Fig. 3e.

Next, we categorized all interactions as direct-stable, direct-unstable, indirect-stable, or indirect-unstable based on multimer prediction scores and MS coverage. All interactions with a score above 0.6 were considered potential direct interactions (represented in blue and green sectors). All proteins in the MS with coverage equal to or greater than that of the TREX2 subunits were categorized as stable interactions. If interacting partners showed both high coverage in MS and a high AF score, we classified them as direct interactions within a stable complex (direct-stable, green sector). Interacting partners that stably associate with the complex but do not interact directly (AF score < 0.6) were placed in the bottom-right purple sector (indirect-stable); all interactors with AF score > 0.6 but low MS coverage were labelled as direct-unstable. All indirect-unstable interactions were located in the bottom left corner and excluded from further analysis (Fig.3e).

### Protein network analysis

All direct-stable, direct-unstable, and indirect-stable interactors were included in the further analysis using the STRING database (version 12.0) (45). Candidates identified from IP-MS were assigned Gene Ontology (GO) terms for specific biological processes and grouped using k-means clustering. Circle sizes and colors were adjusted based on AF scores and GO terms, grouping processes with similar characteristics by color. Colors were assigned to the protein nodes using a Python script supplied by the STRING database, which we subsequently modified to adjust node circle sizes.

All interacting partners from IP-MS were analyzed using several databases, including BioGRID, STRING, and Human Protein Atlas (HPA), as well as a manual literature overview, and categorized to novel and known interactions, which are provided in Supplementary Table 4.

The clustering of IP-MS samples for Figure 3f was performed using Kendall Averaging in Morpheus, https://software.broadinstitute.org/morpheus.

Phylogenetic analysis of Sac3 and Thp1 homologs was performed using SnapGene (version 7.2) and visualized in Geneious Prime version 2023.2.1.

### Degreasing coverslips

Coverslips for immunofluorescence or RNA Fluorescent in situ Hybridization were washed with the solution contained 50% of 10x Tris/Glycine/SDS buffer (BioRad, 161-0732), 25% ethanol, and 25% 1 N NaOH for 5 min to remove residual oils. Subsequently, coverslips were rinsed with deionized water for 15 min, 100% EtOH for 5 min, and dried.

### RNA Fluorescent in situ Hybridization (FISH)

DLD-1 cells were seeded onto degreased coverslips in 12-well plates at a density of 0.1 x 10^6^ cells per well, 2 days before the experiment. The cells were incubated in fresh media with or without 1 mM auxin for 8 h. Afterwards, coverslips were washed with PBS and fixed in 4% PFA at RT for 10 min. The PFA was then aspirated, and the cells were treated with 100% methanol (cooled to -20 °C) for 10 min. Methanol was aspirated and replaced with 70% ethanol for an additional 10 minutes. The ethanol was then removed and replaced with 1M Tris-HCl, pH 8.0, for 5 min. After the Tris-HCl solution was aspirated, the coverslips were treated with a hybridization buffer containing 2x SSC (ThermoFisher, AM9770), 25% formamide, 10% dextran sulfate, 0.005% BSA, 1 mg/ml yeast tRNA (ThermoFisher, 15401029), and 0.1 μM AminoC6-Alexa647-oligo-dT30. The coverslips with cells were incubated with the hybridization buffer for 2 hours at 37 °C in a dark chamber, followed by washes with 4x SSC and 2x SSC. To visualize HA- and FLAG- tagged proteins primary anti-HA (Sigma, 11867423001) or anti-FLAG M2 (Sigma, F1804) antibodies were diluted in 2x SSC buffer, containing 0.1% Triton X100. Coverslips were incubated with antibodies for 1h, then washed 3 times with 2x SCC, and incubated with secondary antibodies diluted in 2x SSC, containing 0.1% Triton X100 for 1h. After staining, coverslips were washed 2x with 2x SCC. To visualize nuclei, cells were stained with 1.5 μg/ml Hoechst 33258 (ThermoFisher, H3569) for 10 min, washed with PBS, and finally mounted in ProLong Gold antifade reagent (ThermoFisher, P36930).

Images were captured as described in the time-lapse fluorescent microscopy method section. A series of 10 z-planes, each 0.2 μm apart, was captured with a 100x ApoTIRF lens, 640-nm laser power at 4.5%, an exposure time of 100 ms, and no binning. Nucleus-to-cytosol (Nu/Cyt) ratio of polyA RNA was quantified in Image J (National Institutes of Health) and plotted with scatter plot function in Prism 10 software. Data are expressed as mean value with standard deviations as error bars. Unpaired two-tailed Student *t*-test has been applied for comparison of untreated and auxin-treated cells.

### Time-lapse fluorescent microscopy

Cells were grown on 4-well glass bottom chambers (Ibidi), imaged on the Eclipse Ti2 inverted microscope (Nikon), equipped with a confocal scanner unit (Yokogawa CSU-W1), PInanoZ/TokaiHit stage top incubator, and controlled by NIS-Elements AR 5.21.03 software (Nikon) utilizing Nikon CFI Apochromat TIRF 60XC Oil and Plan Fluor 40x/1.30 Oil immersion objective lens. Cells were imaged in FluoroBrite DMEM (ThermoFisher, A18967-01) media. The microscope was equipped with temperature-, CO_2_- and humidity-controlled chamber that maintained a 5% CO_2_ atmosphere and 37°C. NG and mCherry fluorescent protein signals were excited with a 488-nm (no more than 20% of power was applied) and 568-nm (no more than 50% of power was applied) laser lines, respectively. A single image consisting of a series of 0.5 µm optical sections was acquired. Images analyzed using Fiji software (National Institutes of Health, version 2.16.0/1.54p). Images represent maximum intensity projections of entire z-stacks or a single z-stack.

### Immunofluorescence staining

Untagged DLD-1 cells or DLD-1 expressing endogenously tagged PCID2, LENG8 or SAC3D1 were seeded on 12 mm #1 coverslips (Electron Microscopy Sciences) at a density of 1.2*10^5^ cells on 12-well plates and grown for 2 days. PCID2 depletion was achieved using 3-indoleacetic acid (auxin, IAA, 1:500) (Sigma, I3750-5G-A) for 4h. LENG8 or SAC3D1 depletion was achieved using 5-Ph-IAA (1:1000) (MCE, HY-134653) for 4h. The control cells were either untreated (PCID2) or treated with DMSO (LENG8, SAC3D1). Cells were washed with PBS (pH 7.4) and fixed under the following conditions: 4% paraformaldehyde (Electron Microscopy Sciences, 15710) containing 0.5% Triton X-100 in PBS for 15 min at room temperature (RT) (Figure 6h; Supplementary Fig. 6b-c; Supplementary Fig. 7d-e); 4% paraformaldehyde alone (Supplementary Fig. 6b-c); or methanol alone (Supplementary Fig. 6b-c). For nuclear-enriched images (Supplementary Fig.7b-c), cells were prewashed for 5 min at RT with buffer containing 20 mM HEPES (pH 7.8), 2 mM DTT, 10% sucrose, 5 mM MgCl₂, 5 mM EGTA, 1% Triton X-100, and 0.075% SDS. In all cases, coverslips were subsequently blocked with 10% horse serum (Vector Laboratories, S-200-20) for 1 h. Intracellular protein localization was detected using specific primary antibodies and Alexa Fluor–conjugated secondary antibodies. Nuclei were visualized with 1.5 μg/ml Hoechst 33258 (ThermoFisher, H3569). Coverslips were mounted in ProLong Gold antifade reagent (ThermoFisher, P36930).

Images were acquired as described in the Time-Lapse Fluorescence Microscopy section. Brightness and contrast adjustments were applied equally to all images within each experiment using Fiji software. RGB images generated in Fiji were further processed using Adobe Photoshop (version 27.2.0) and Adobe Illustrator (version 30.1).

### Antibodies

The following antibodies were used for immunostaining, immunoprecipitation, and Western blot analysis: anti-rabbit (ThermoFisher, A11034) AlexaFluor-488 conjugated antibodies (Invitrogen); anti-mouse (ThermoFisher, A11004) AlexaFluor-568 conjugated antibodies (Invitrogen). Specific primary antibodies against HA (Sigma, 11867423001), FLAG M2 (Sigma, F1804), ACTB (Cell Signaling, 13E5, #4970), GANP (Abcam, ab113295), PCID2 (ab216042 IP only), RFP (Chromotek, 6G6), ENY2 (ThermoFisher, MA5-27843), CETN2 (ProteinTech, 15877-I-AP), CETN3 (ProteinTech, 15811-I-AP), LENG8 (Fortis, A304-947A), SAC3D1 (ProteinTech, 25857-I-AP), DDX39B (ProteinTech, 14798-I-AP), PNN (ProteinTech, 18266-I-AP), EIF3I (ProteinTech, 11287-I-AP), SNRNP200 (ProteinTech, 23875-I-AP), SRPK1 (ProteinTech, 14073-I-AP), and SC35 (Abcam, ab11826). Secondary HRP-conjugated anti-mouse and anti-rabbit antibodies for Western blot analysis were purchased from Sigma-Aldrich.

### RNA-seq samples preparation

AID-targeted cells at 3^rd^-6^th^ passage were seeded on 6-well dishes. One million of adherent DLD-1 cells were washed with PBS and lysed directly with RLT buffer from RNAeasy Plus Mini Kit (Qiagen, 74134). Extracted RNA was analyzed on Qubit 4 Fluorometer (ThermoFisher) with Qubit RNA BR Assay Kit (ThermoFisher, Q10211), Qubit RNA IQ Assay Kit (ThermoFisher, Q33222), and Bioanalyzer 2100 (Agilent). Samples with RIN value more than 9 were used for the library construction. GEO accession GSE291063 has two separate sets of data. RNA samples purified in 2021 were depleted for ribosomal RNA before library construction using Ribo-Zero™ Gold Kit H/M/R Kit (Illumina). RNA samples purified in 2024 were enriched for poly (A) RNA as part of the TruSeq stranded kit workflow using total RNA as input. PolyA-enriched RNA-Seq libraries were constructed using the Illumina TruSeq Stranded mRNA Library Prep Kit according to the manufacturer’s instructions with specific barcodes. All the RNA-Seq libraries were pooled together and sequenced using Illumina NovaSeq 6000 to generate approximately 40 million 2×100 paired-end reads for each sample. The raw data were demultiplexed and analyzed further.

### RNA-seq Analysis

Raw RNA-Seq data reads were trimmed of adapters with Cutadapt (version 4.4)(46) with the parameters ‘-a AGATCGGAAGAGCACACGTCTGAACTCCAGTCA -AAGATCGGAAGAGCGTCGTGTAGGGAAAGAGTGT —nextseq-trim 20 --overlap 6 --minimum-length 25’. Alignment of trimmed reads was performed with HiSat2 (version 2.2.1)(47) with default parameters against the GENCODE GRCh38 human genome build. Transcript-based read quantitation was performed using Kallisto (version 0.48.0)(48). Read counts were aggregated to genes with the R package tximport (version 1.26.0). Read counts were normalized by Relative Log Expression (RLE) method (49) and evaluated for differential expression using the Bioconductor package DESeq2 (version 1.38.0) (50), comparing defined sample sets as biological replicates and drawing pairwise comparisons. Differentially expressed genes were clustered in heatmaps using the R package Mclust (v6.1.1) (51). Enrichment analysis of subsets of altered genes was performed using the Bioconductor package clusterProfiler (version 4.6.0) (52).

Differential exon and intron usage analysis was performed using DEXseq (version 1.52.0) (53, 54) on exon and intron read counts generated with SubRead featureCounts (version 2.0.3). The exon and intron annotation was prepared with the functions exonicParts and intronicParts respectively from the R package GenomicFeatures (version 1.50.2). Heatmaps represent the combined fraction of differentially used exons and introns, calculated as the number of significant exons and introns normalized to the total number of exons and introns, respectively, for each strain. Only genes with significant exons and/or introns in at least 1 strain are represented. To plot the heatmap, we utilized the same function developed for the DE analysis clustered heatmap. The Venn diagrams were created using Venny 2.1 tool (55) and DeepVenn (56). Gene Ontology (GO-terms) enrichment analysis was performed using an integrated HumanMine database (v12 Feb 2022) (57) with Holm-Bonferroni test correction (p-value 0.05-0.1) for Biological Processes. Reads from bigwig files were visualized in IGV (version 2.17.0) against primary assembly from GENCODE v28 (GRCh38.p12). Location of polyA sites were determined using PolyASite 2.0 (58) (Supplementary Table 8). GTEx Portal was used to determine abundance of transcripts encoding subunits of the TREX2 complex. The data in Supplementary Figure 1 were obtained from the <u>GTEx Portal</u> and <u>dbGaP</u> accession number phs000424.v10.p2 on 12/22/2025, using GTEx Multi Gene Query function.

### RT-PCR and RT-qPCR

A total of 3 μg of RNA was purified according to the protocol described in the RNA-seq sample preparation section and reverse transcribed into cDNA using the SuperScript IV kit (Thermo Fisher Scientific, 18090050) in the presence of oligo(dT)18 primer and RNaseOUT Recombinant RNase Inhibitor, as recommended by the manufacturer. Quantitative PCR was performed using iTaq Universal SYBR Green Supermix (Bio-Rad, 172-5120) on a CFX Connect™ Real-Time PCR Detection System (Bio-Rad). To analyze changes in differential exon and intron usage events, region-specific primers were designed. A dilution series was used to generate standard curves for all primer sets. Only primer sets with amplification efficiency (E) between 85% and 115%, an R² value > 0.9, and a slope between −3.58 and −3.10 were used for subsequent quantification. Dilution series and Ct values were analyzed using Bio-Rad CFX Manager 3.1 (version 3.1.1517.0823). To quantify fold changes, the upregulated region was normalized to an upstream or downstream region of the same transcript that showed no change. Changes in the upregulated region were also compared with corresponding changes observed in auxin-treated DLD-1 cells. Primer sets used for RT-qPCR and RT-PCR are listed in Supplementary Table 1. Relative abundance of transcript isoforms was plotted using a scatter bar plot function in Prism (version 10.2.3). Data are presented as mean ± SD. Statistical significance between untreated and auxin-treated cells was assessed using an unpaired two-tailed Student’s t-test.

---

**Figure S1.**
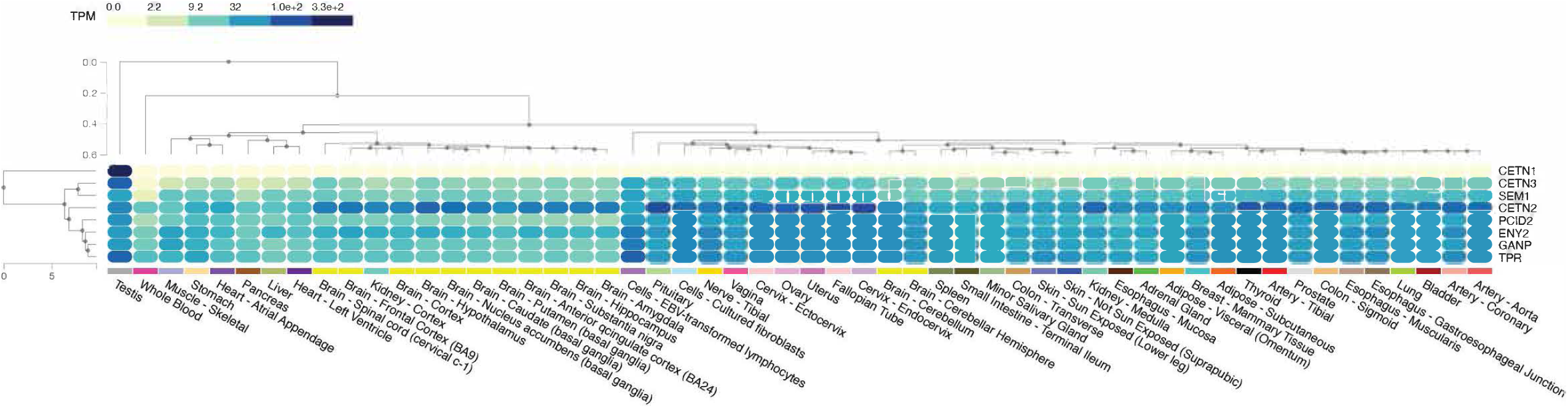
TREX2 subunits differentially expressed across human tissues. GTEx heat map of TREX2 subunits and TPR RNA transcripts abundancy across 54 human tissues. The heat map is ordered by tissue clustering. A gradient from light yellow to dark blue represents abundance of transcripts across tissues.

**Figure S2.**
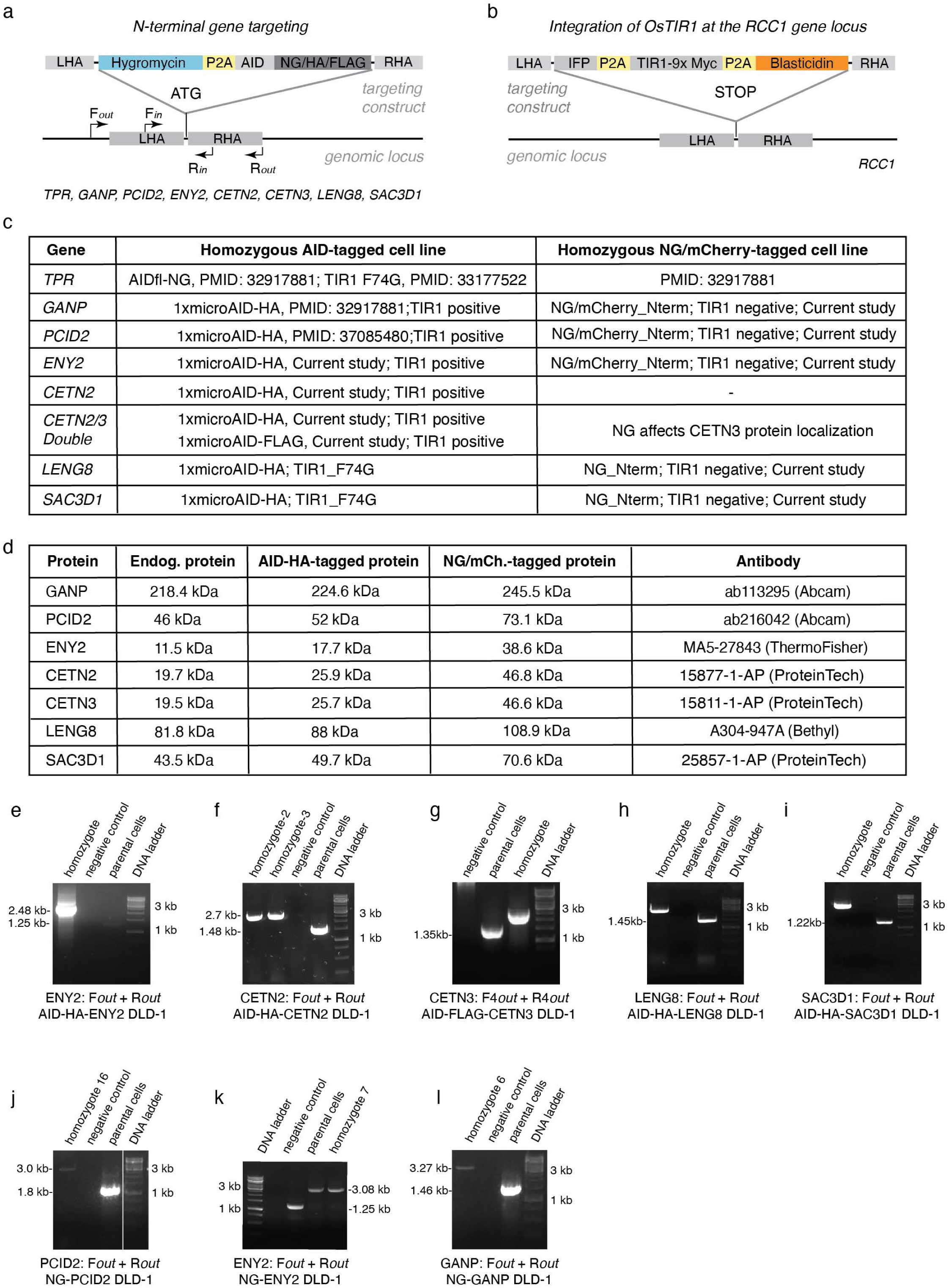
TREX2 subunits, LENG8 and SAC3D1 gene targeting strategy. **a,** A scheme of endogenous targeting of TREX2 subunits and TREX2 subunits’ associated protein partners with hygromycin-P2A-AID-NG/HA/FLAG tags. A position of two primer sets used for genotyping is shown relative to homologous arms. All genes were N-terminally targeted. F*out*-R*out* primer set anneals outside of the homologous arms, F*in*- R*in* primer set anneals inside of the homologous arms. **b**, A scheme of *Oryza sativa* TIR1 genomic integration into the *RCC1* gene locus. Two alternative types of TIR1 and auxin have been used in this study: a canonical OsTIR1 along with 1 mM Indole-3-acetic acid sodium salt and OsTIR1_F74G mutant along with 1 uM 5-Ph-IAA (42). **c**, A list of cell lines used in this study. **d**, Molecular weight of endogenous and HA- or NG/mCherry- tagged TREX2 subunits and TREX2 subunits’ associated protein partners. Last column shows commercial antibody used to detect proteins on Western blot to confirm targeting and degradation. **e-i**, Genomic PCR of homozygous clones demonstrating the integration of the Hygromycin-P2A-AID-HA/FLAG sequence into genomic loci of ENY2 (e), CETN2 (f), CETN3 (g), LENG8 (h), and SAC3D1 (i) genes with *out-out* primer set. AID – Auxin Inducible Degron, HA - human influenza hemagglutinin. **j-l**, Genomic PCR of homozygous clones demonstrating the integration of the Hygromycin-P2A-NG sequence into genomic loci of PCID2 (j), ENY2 (k), and GANP (l) genes. NG – Neon Green fluorescent protein. NG-tagged cell lines (j-l) were used for IP-MS experiment, see Figure 3.

**Figure S3.**
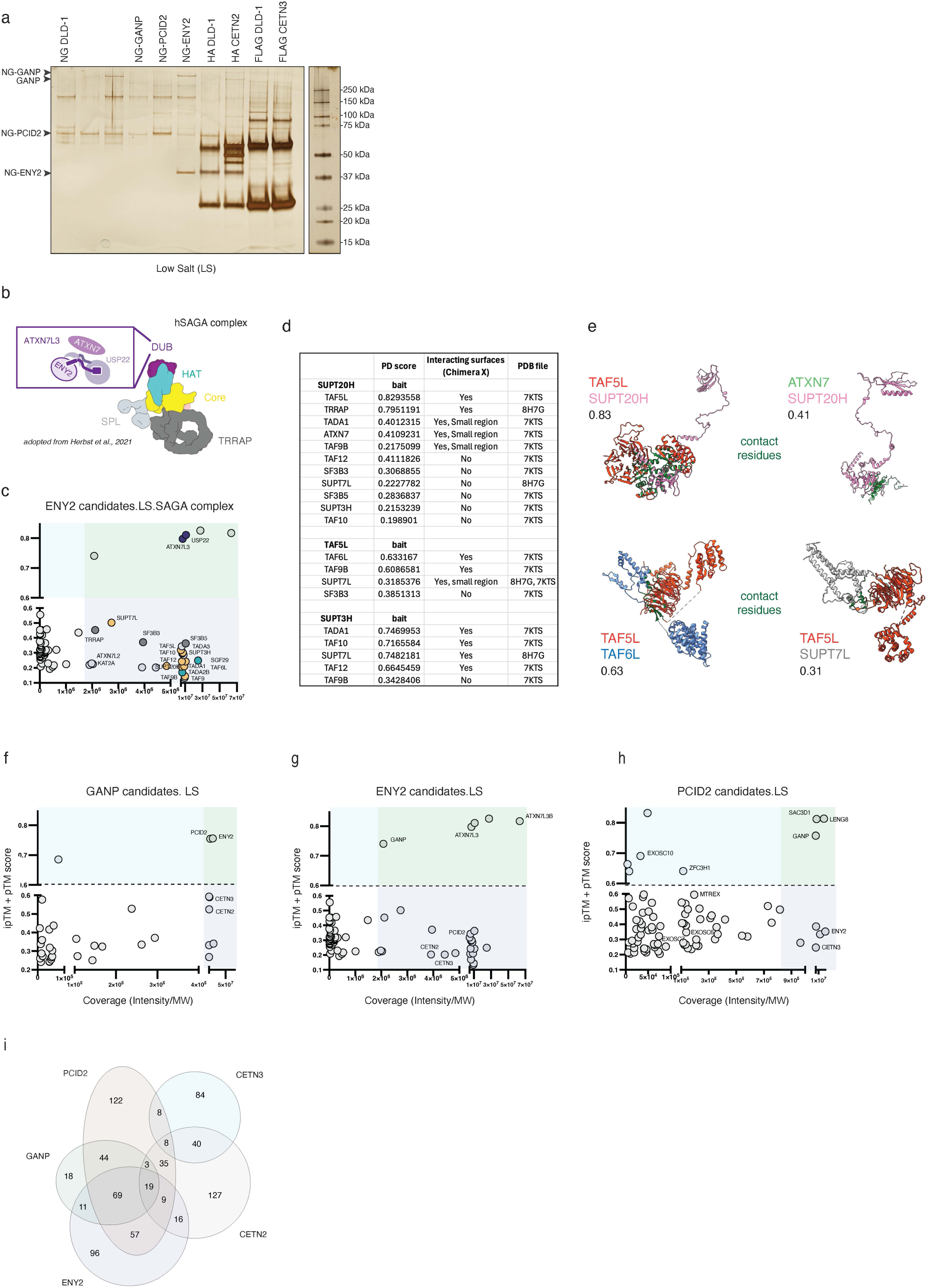
Binding partners of the TREX2 subunits under LS purification condition and *in silico* AF screen of the SAGA complex subunits. **a**, A silver gel of NG-GANP, NG-PCID2, NG-ENY2, HA-CETN2, and FLAG-CETN3 and associated proteins isolated by the immunoprecipitation using the low salt buffer and NG-, HA, or FLAG-beads, respectively. The DLD-1 cell line was used as a control to assess background binding of the beads. NG-GANP, NG-PCID2, and NG-ENY2 are indicated with arrows. **b**, A schematic of SAGA complex domains. **c**, A quartile plot of ENY2 protein partners highlighting SAGA complex subunits immunoprecipitated with NG-ENY2. The right axis shows the protein coverage in the MS divided by its molecular weight. The left axis indicates the sum of the predicted Template Modeling score (pTM) and interface pTM score (ipTM), which are measures of predicted structure accuracy generated by AF for the protein and protein-protein binding interphase. The score 0 indicates no interaction between the bait and the prey; the score 1 indicates a high chance of both proteins interacting. DUB, HAT, SPL, Core, and TRRAP domain subunits are colored in violet, cerulean, light grey, yellow, dark grey colors, respectively on graph and schematic. **d**, *In silico* AF screen of known SAGA complex subunits interacting partners. Interacting surfaces were determined in Chimera X based on Cryo-EM PDB files. **e**, The visualization of interacting residues of selected SAGA subunits in Chimera X. The structures are derived from the 3D atomic structures of the cryo-EM density maps of 7KTS or 8H7G. The corresponding AF scores of the multimers are shown in the table, above or below each example. **f-h**, A quartile scatter plots of GANP (f), ENY2 (g), and PCID2 (h) binding partners under low salt (LS) purification condition. **i**, A Venn diagram representing the number of overlapping targets among binding partners of GANP, PCID2, ENY2, CETN2, or CETN3.

**Figure S4.**
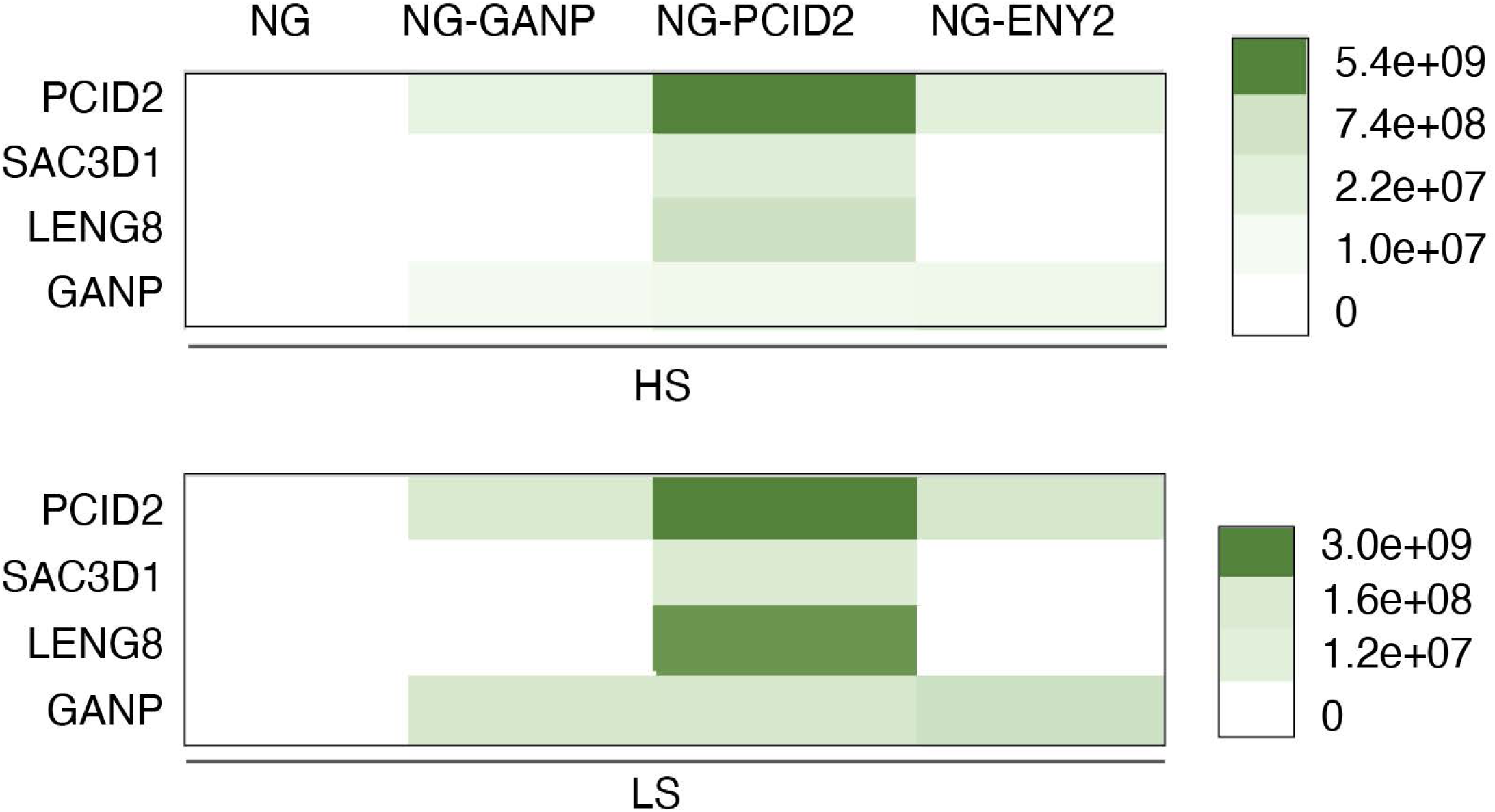
Identification of novel PCID2 interacting partners. A heat map comparing the MS abundance (in green) of PCID2 binding partners across the NG-GANP, NG-PCID2, and NG-ENY2 cell lines.

**Figure S5.**
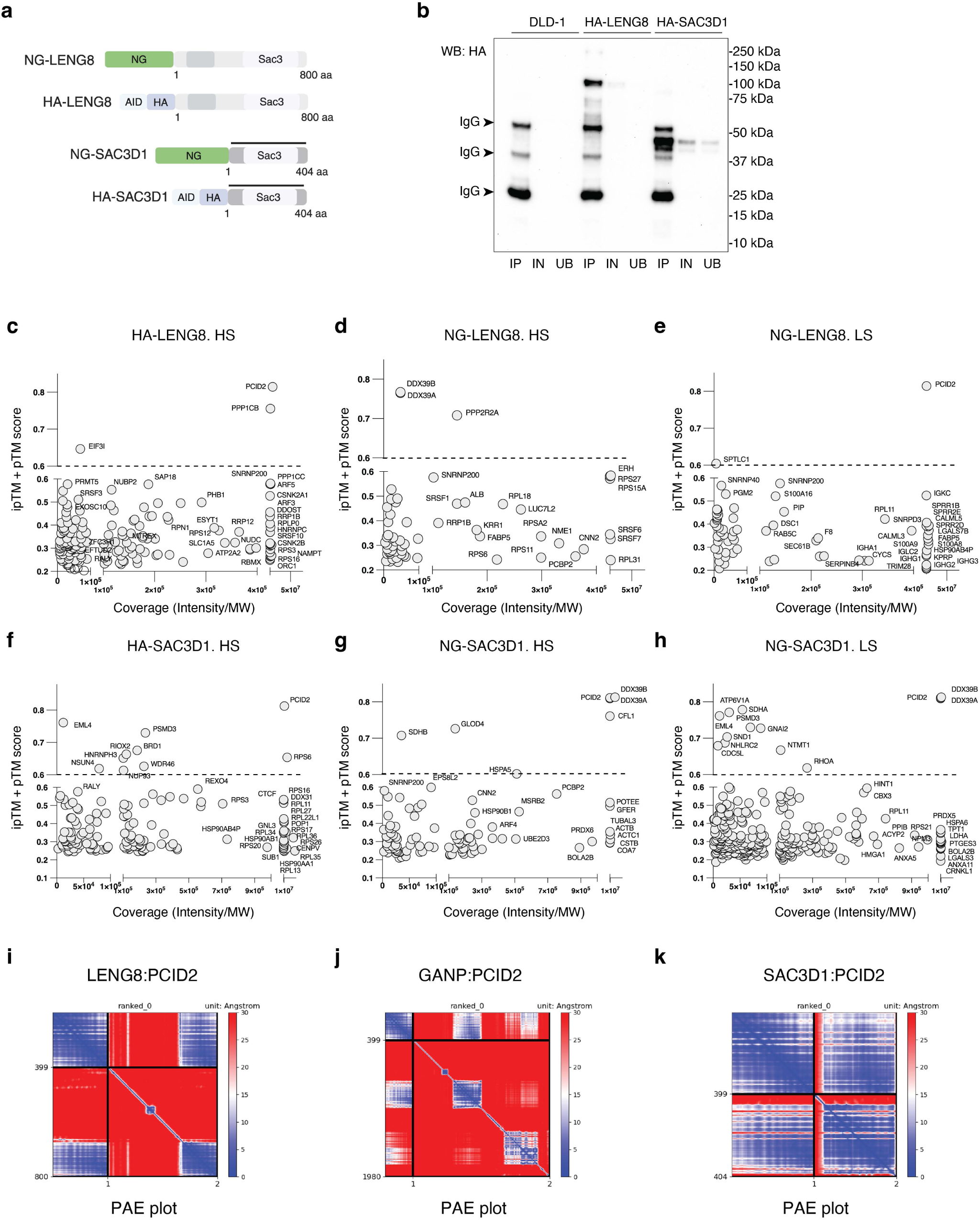
Identification of LENG8 and SAC3D1 interacting partners. **a**, A scheme of endogenous targeting of LENG8 and SAC3D1 with NG or AID-HA tags. **b**, An example of western blotting showing precipitation of HA-tagged LENG8 and SAC3D1 from DLD-1, AID-HA-LENG8, and AID-HA-SAC3D1 cell lines under high salt extraction conditions. **c-h**, Quartile plots showing LENG8 and SAC3D1 binding partners under high (HS) and low salt (LS) purification conditions. **i-k**, Predicted aligned error (PAE) plots of LENG8:PCID2, GANP:PCID2, and SAC3D1:PCID2 multimers.

**Figure S6.**
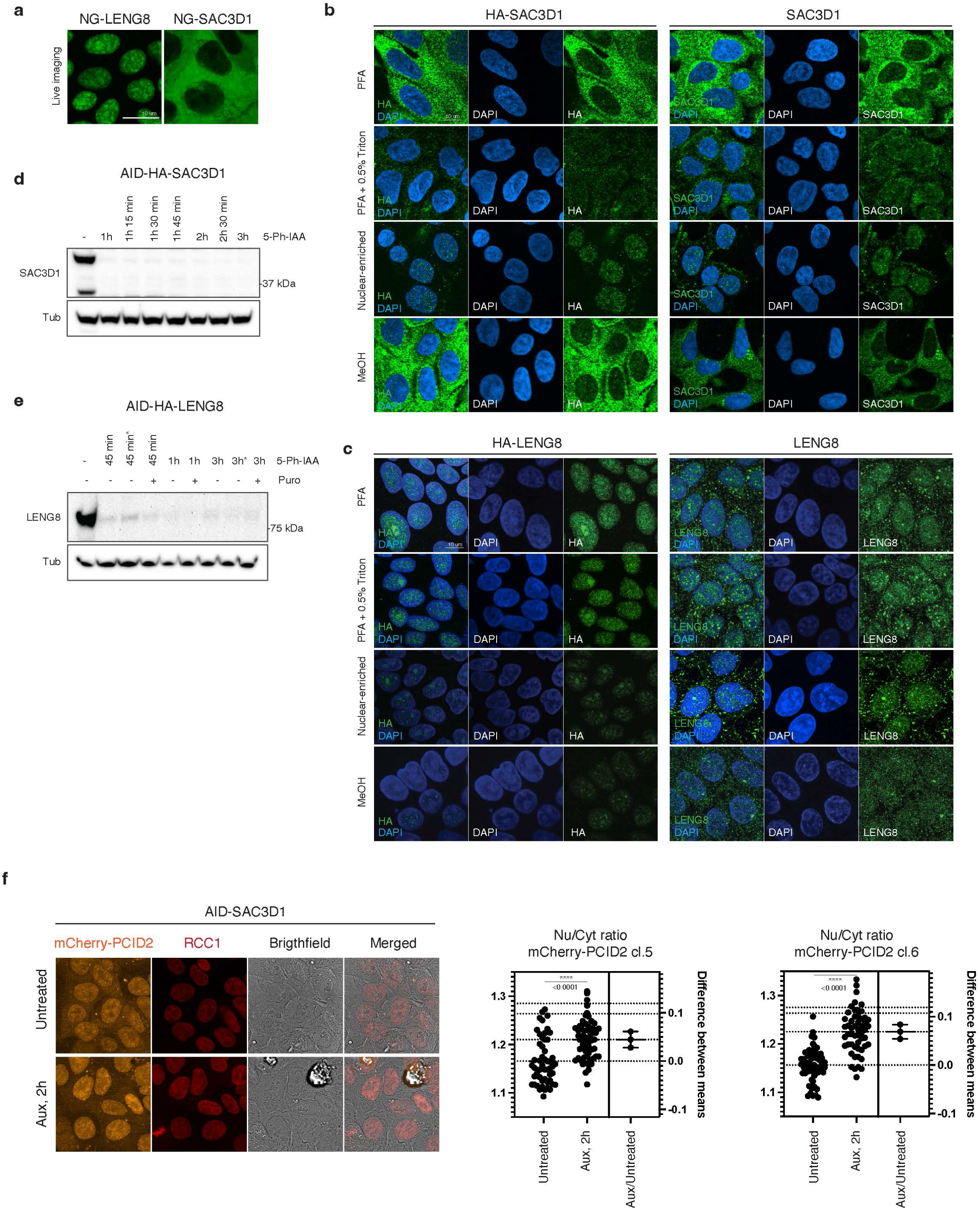
Intracellular localization of endogenous and tagged LENG8 and SAC3D1. **a**, Live imaging of NG-tagged LENG8 and SAC3D1 in DLD-1 cell line. **b-c**, Intracellular localization of HA-tagged and untagged SAC3D1 (b) and LENG8 (c) using four alternative fixation condition methods. PFA – 4% Paraformaldehyde; PFA + 0.5% Triton – cells permeabilized simultaneously; nuclear-enriched – cells washed with buffer A (20 mM HEPES, pH 7.8; 2 mM DTT, 10% sucrose, 5 mM MgCl_2_, 5 mM EGTA, 1% TX100, 0.075% SDS) for 5 min and fixed with 4% PFA; MeOH – cells fixed and permeabilized with methanol. **d**-**e**, Western blotting of untreated or 5-Ph-IAA treated AID-HA-SAC3D1 (d) and AID-HA-LENG8 (e) cell lines. Note the degradation of SAC3D1 and LENG8 proteins happens within 1 h and 45 min, respectively. *Indicate replacement of culture media with fresh media containing 5-Ph-IAA every 30 min. **f**, A single plane image and estimation plots showing the difference between means in the nuclear-to-cytoplasmic (Nu/Cyt) localization ratio of mCherry-PCID2 in untreated cells compared to those with an acute loss of SAC3D1. Data are presented as mean values, ****p-value < 0.0001, unpaired Student’s *t*-test. Two separate clones were analyzed: clone 5, n = 52 untreated and n = 59 5-Ph-IAA-treated cells; clone 6, n = 57 untreated and n = 53 5-Ph-IAA-treated cells.

**Figure S7.**
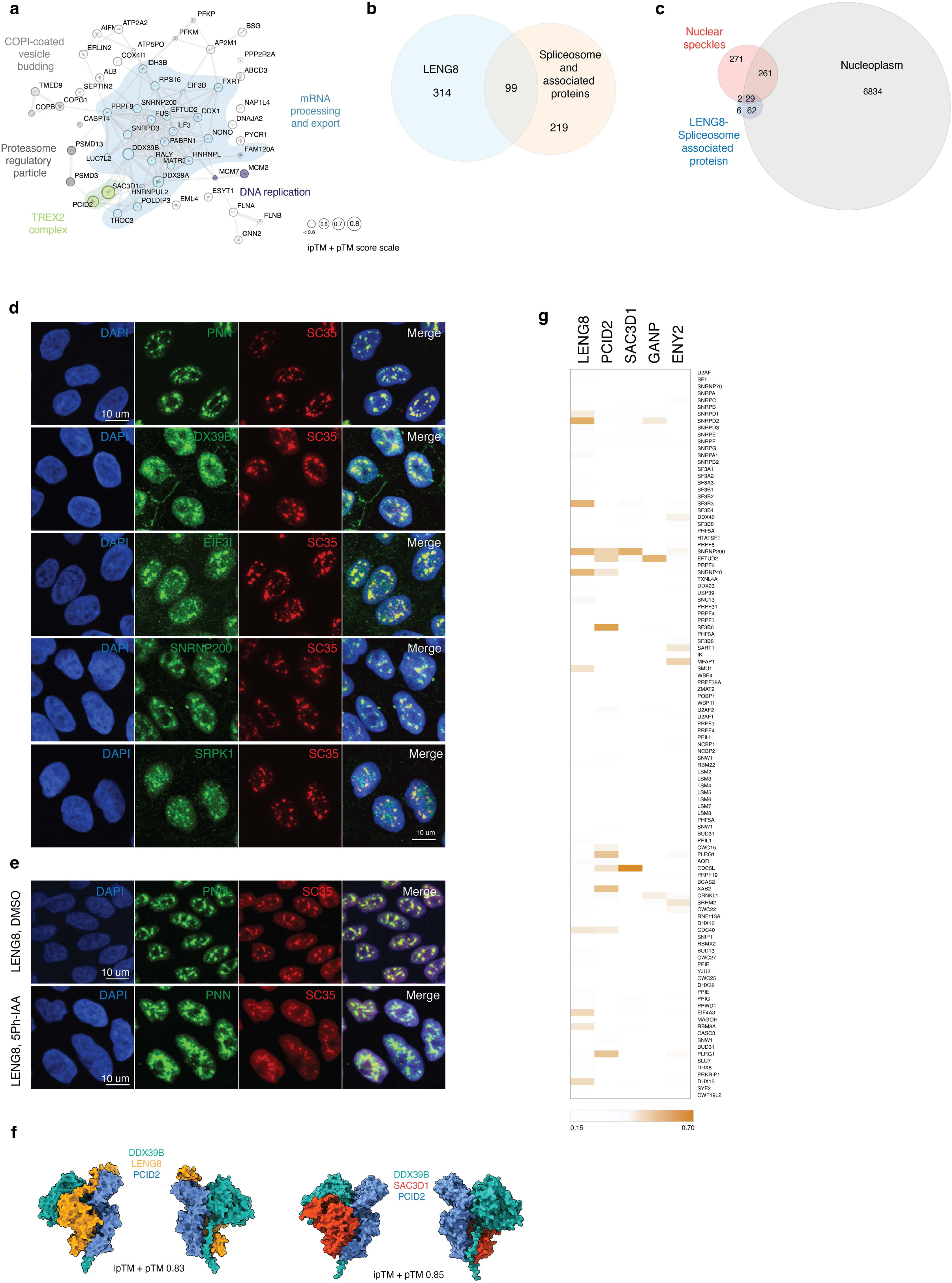
LENG8-bound proteins enriched in nuclear speckles. **a**, STRING map of SAC3D1 interacting partners reproduced in at least two independent purification conditions with 5-fold enrichment. Circles corresponding to proteins with AF scores 0.6 and higher are enlarged respectively. **b**, A Venn diagram of LENG8 binding partners showing the overlap with spliceosome and spliceosome-associated proteins. **c**, A Venn diagram illustrating the association of LENG8-spliceosome-associated proteins (99 candidates from b) with the pool of nucleoplasmic proteins and proteins localized in nuclear speckles. **d-e**, Co-localization of nuclear speckle marker SC35 and top LENG8 interacting partners in untreated DLD-1 cells and upon LENG8 loss (**e**). **f**, AF multimers of PCID2 protein with LENG8 or SAC3D1, and DDX39B. **g**, A heat map of AF prediction scores for LENG8, PCID2, SAC3D1, GANP, ENY2 and spliceosome components.

**Figure S8.**
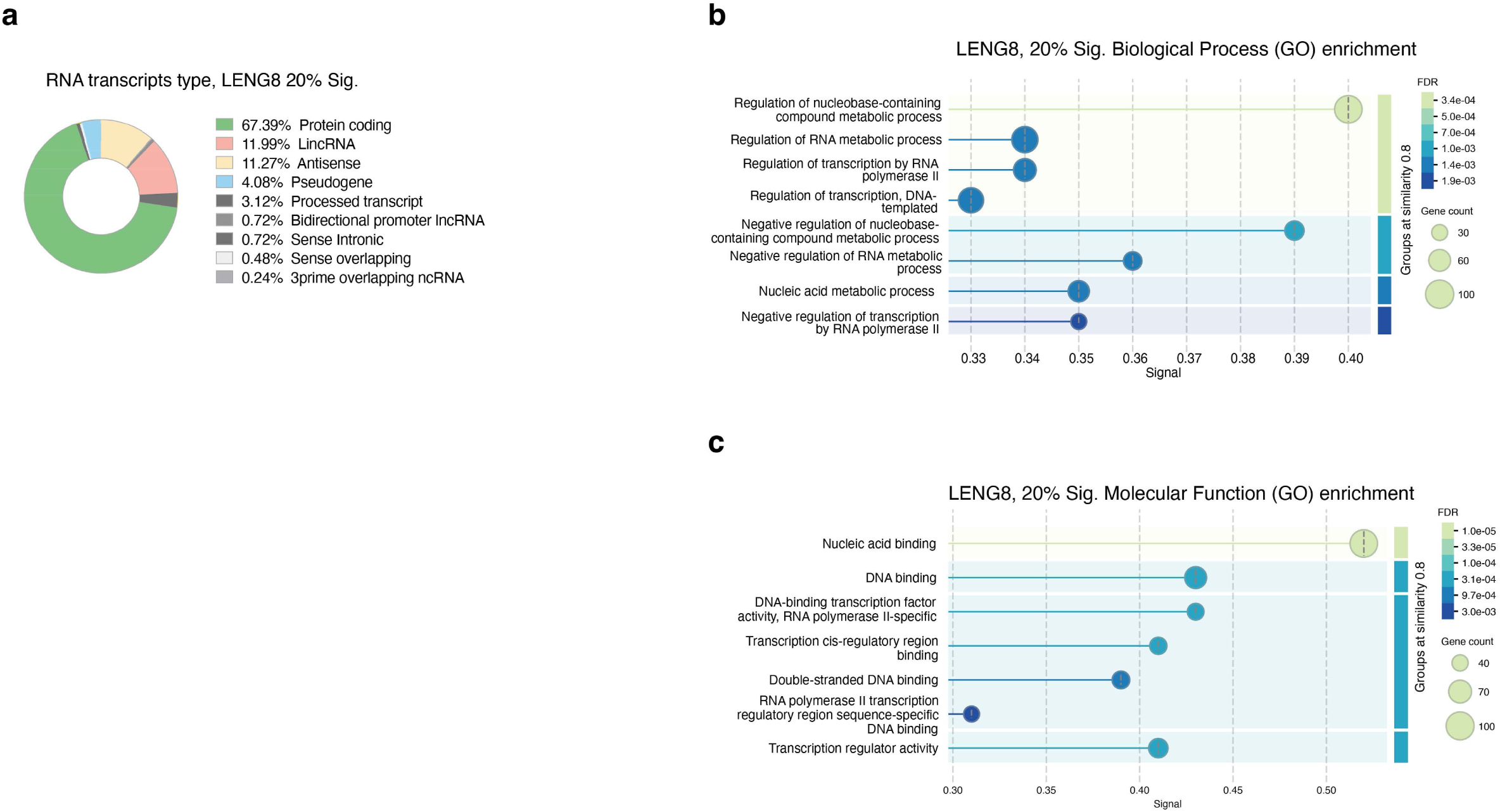
Gene expression changes upon LENG8 loss. **a**, A proportion of RNA transcript subtypes in LENG8 RNA-seq data. Only significantly changed differentially expressed RNAs were taken into analysis with log2FC > 20%, adj. p-value < 0.05. **b**-**c**, Top GO-terms (Biological Process, **b**; Molecular Function, **c**) of differentially expressed RNAs upon LENG8 loss (HumanMine v12, 2022).

**Figure S9.**
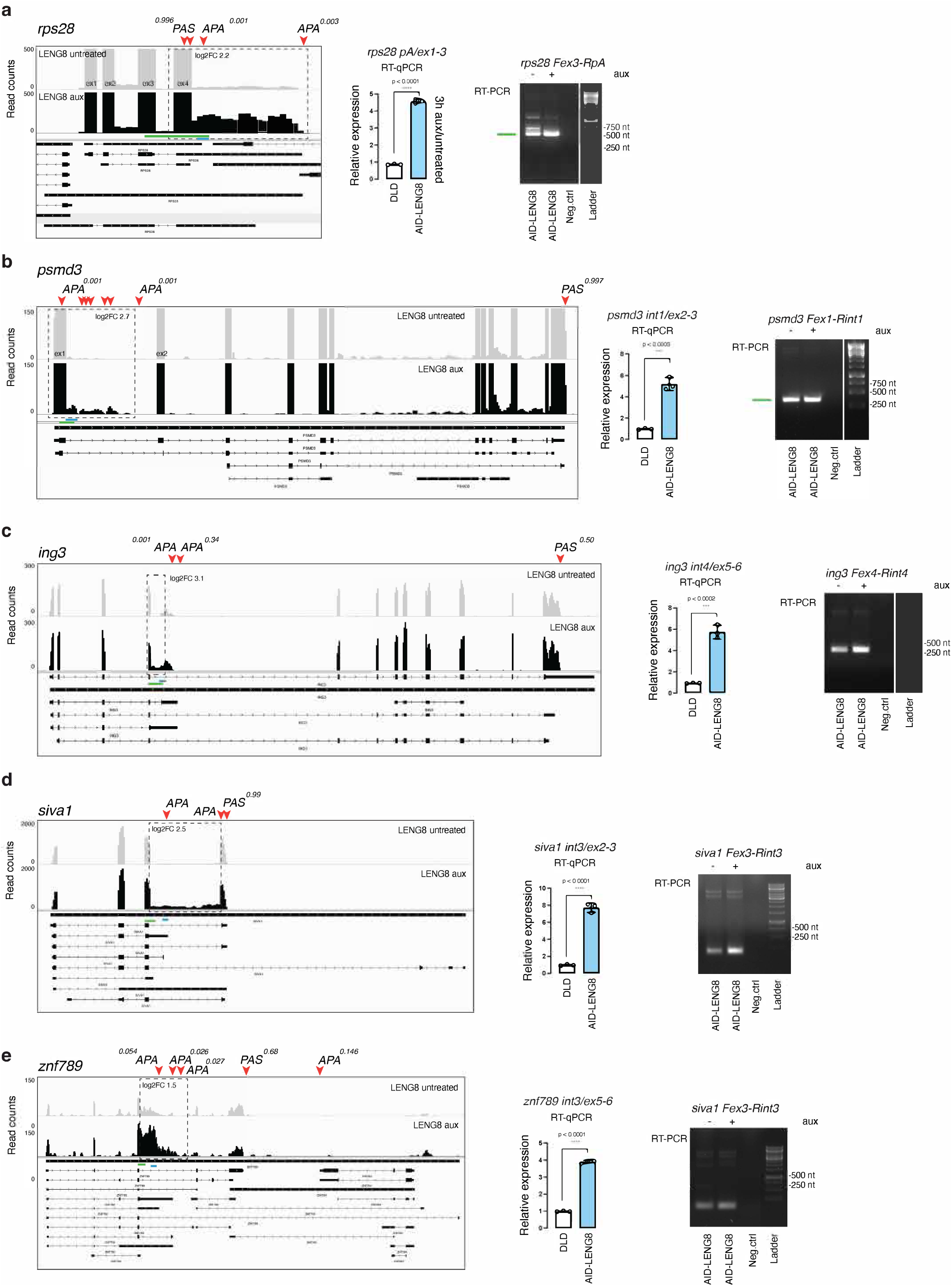
Examples of alternative polyadenylation site usage events upon LENG8 loss. **a**-**f**, IGV snapshots of RNA-Seq data, RT-qPCR analysis of upregulated regions (blue line), and agarose gel electrophoresis of the RT-qPCR products between the last exon and the joining upregulated intron (green line) for *rps28* (**a**), *psmd3* (**b**), *ing3* (**c**), *siva1* (**d**), and *znf789* (**e**) transcripts. Regions with alternative polyadenylation usage events with increased read counts are highlighted with dashed squares. The frequency of PAS usage and fold change of the upregulated region are indicated, respectively. RT-qPCR graphs represent the mean value of three technical replicates of one experiment; error bars are SD. Asterisks indicate ****p-value < 0.0001 and ***< 0.01 in unpaired two-tailed Student’s *t*-test.

